# Photosynthetic acclimation mediates exponential growth of a desert plant in Death Valley summer

**DOI:** 10.1101/2023.06.23.546155

**Authors:** Karine Prado, Bo Xue, Jennifer E. Johnson, Sterling Field, Matt Stata, Charles L. Hawkins, Ru-Ching Hsia, Hongbing Liu, Shifeng Cheng, Seung Y. Rhee

## Abstract

Heat waves, now more frequent and longer due to climate change, devastate plant productivity. Although rare, thermophilic plants could hold keys to engineering heat resilience in crop plants. *Tidestromia oblongifolia* is a thermophilic flowering plant that thrives at temperatures above 45°C. When exposed to Death Valley summer conditions, *T. oblongifolia* increased its thermal optimum of photosynthesis within a day and accelerated growth within 10 days. The physiological changes were accompanied by morphological, anatomical, and gene expression changes revealed by a newly sequenced genome. In bundle sheath cells where Rubisco fixes CO_2_, mitochondria relocated to chloroplasts and novel, cup-shaped chloroplasts appeared. Understanding how this plant acclimates under heat may afford new ways of engineering heat tolerance in crop plants.

**One-Sentence Summary:** *Tidestromia oblongifolia*’s acclimation to Death Valley is accompanied by changes in gene expression, organellar dynamics, and photosynthesis.

## Main Text

Current climate models predict a global surface temperature increase of 1.5-5°C by 2100 (*1*), increase of prolonged extreme events (*2*), reduction of crop productivity (*3*), vegetation shifts (*4*), and changes in species distributions (*5*). Heat waves lead to losses of billions of dollars in agricultural outputs (*2*). Major agricultural crops do not tolerate temperatures above 35°C (*6*). Even sorghum, a bioenergy crop considered to be relatively heat tolerant, decreases its biomass and seed yield at temperatures beyond 40°C (*7*). It is therefore pressing to design innovative breeding and engineering strategies to sustain agricultural yields in the face of increasingly severe climate impacts (*8*). However, bioengineering crop resilience to increasing temperature is challenging because heat tolerance traits are multigenic and not well characterized at the molecular and cellular levels, the functional integration of multiple traits is incompletely understood, and increasing thermotolerance traits can trade off for growth and yield (*8, 9*).

Thermophilic plants represent an interesting source of heat adaptive mechanisms for agricultural applications (*10*). *Tidestromia oblongifolia* is a thermophilic plant native to Death Valley, an environment that holds the highest air temperature officially recorded (*11, 12*). *T. oblongifolia* belongs to the Amaranthaceae family, which includes crops like quinoa, amaranth, spinach, and sugar beet (*13*). A perennial highly adapted to high temperature and high light, *T. oblongifolia* has an optimal photosynthetic rate at 47°C *in situ* (*14*). This plant operates a C_4_ carbon assimilation pathway (NADP-ME C_4_) (*15*), which partitions photosynthesis in specialized cells and organelles to concentrate CO_2_ around Rubisco and suppress its oxygenase activity. C_4_ plants typically grow faster than C_3_ plants in warmer and drier climates, colonize harsh environments, and include highly productive and economically valuable crops such as sugarcane and maize (*16, 17*). How *T. oblongifolia* thrives under extreme heat still remains largely uninvestigated at the molecular, cellular, and genetic levels and no genomic information is available for *T. oblongifolia*.

To understand the basis of *T. oblongifolia*’s thermotolerance, we grew the plants at moderate temperatures and studied the physiological, morphological, and anatomical changes it underwent after being transferred to Death Valley summer conditions. We also sequenced the genome and examined transcriptomic changes in response to high temperature and light, which highlighted putative candidate genes for achieving thermotolerance. The resources and findings from this study open many opportunities and directions to both studying thermotolerance mechanisms and applying thermoadaptive traits to crop plants.

### Death Valley summer elicits exponential growth in *T. oblongifolia*

To examine the behavior of *T. oblongifolia* under conditions that were realistic but controlled, we customized growth chambers to recreate the light and temperature regimes that are characteristic of Death Valley in mid-summer (Fig. S1-S2, Data S1). Nine-week-old *Tidestromia oblongifolia* plants were transferred to Death Valley (DV) summer conditions and monitored for 1 month (Fig. 1A, Fig. S3, Data S1). We included *Amaranthus hypochondriacus* in our study as it is the most closely related C_4_ species to *T. oblongifolia* with a sequenced genome (*13*). We compared two accessions of *T. oblongifolia*, Death Valley (DV) and Dos Palmas (DP), to two accessions of *A. hypochondriacus*, India (IN) and Nebraska (NE), with contrasting maximal temperatures in their native regions (Fig. S4, Data S1). DV summer arrested growth of *A. hypochondriacus* (Fig. S5). In contrast, both accessions of *T. oblongifolia* thrived under these conditions with an exponential growth rate that tripled their biomass within 10 days (Fig. 1BC, Data S1). While growth rate and biomass were not significantly distinguishable between the two accessions, we observed a trend toward higher biomass in the DV accession compared to DP accession (Fig. 1C, Fig. S6A, Data S2). New leaves formed under DV summer were smaller and increased trichome density (Fig. S6B, Data S2), which are morphological changes consistent with thermoadaptation described for desert plants (*18, 19*). Overall, *T. oblongifolia* accessions acclimated to these controlled DV summer conditions and grew exponentially within 10 days.

**Fig. 1.**
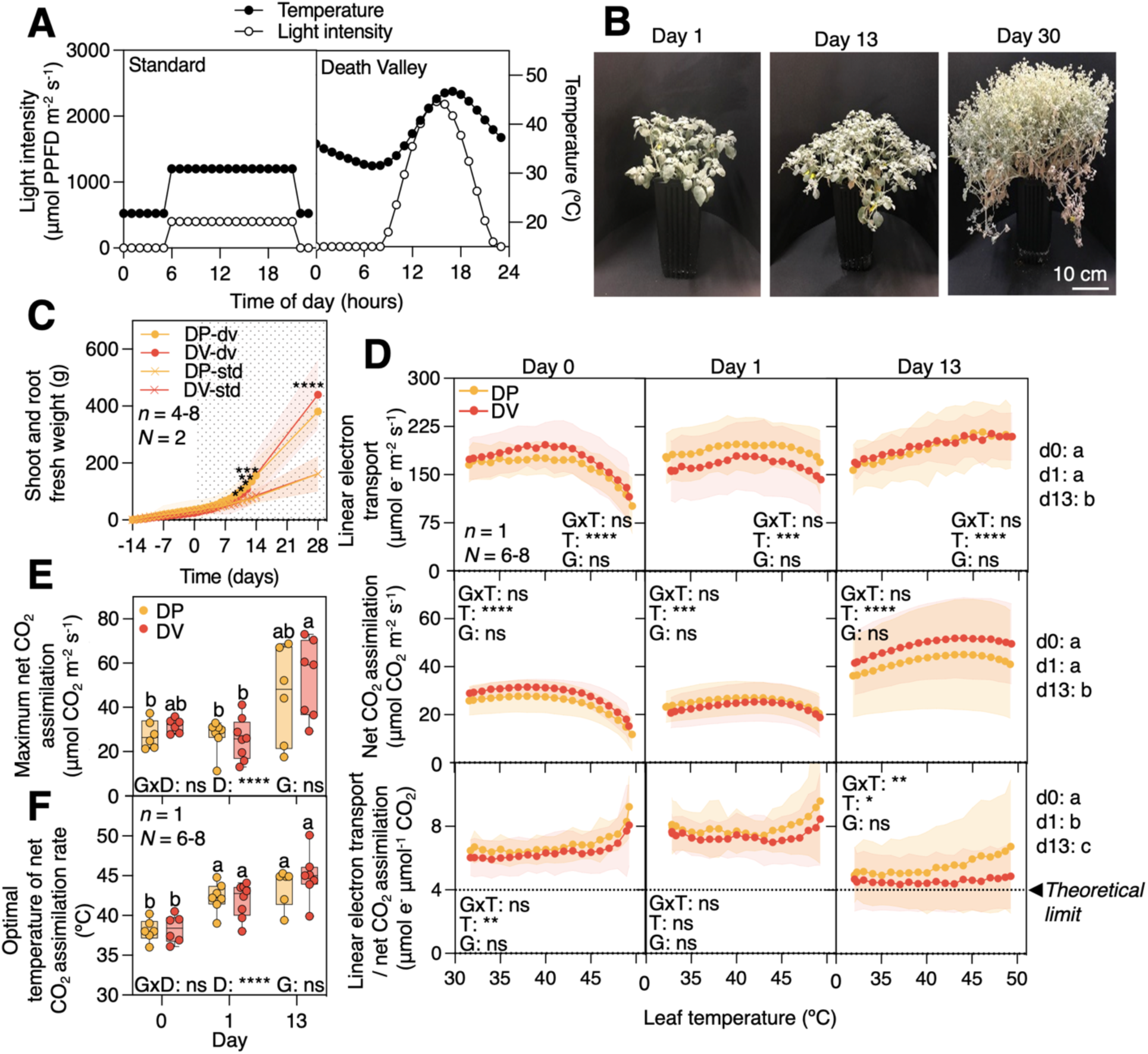
Growth and photosynthetic heat tolerance of *T. oblongifolia* in response to DV summer. (**A**) Daily light and temperature growth conditions. (**B**) Representative images of DV accession challenged with DV summer. (**C**) Average shoot and root fresh weight per plant for 6 weeks for DP and DV accessions. 0 indicates the day prior to the start of treatment under DV summer and the gray area indicates time after transfer. Asterisks indicate statistically significant differences between standard and DV summer conditions using a two-tailed Mann-Whitney test at *P* < 0.05 (*), < 0.01 (**), < 0.001 (***), < 0.0001 (****). (**D**) Thermal response curves of photosynthetic performances. For (C) and (D), circles and filled areas represent means and ±95% confidence intervals. Asterisks inside the boxes indicate statistically significant differences between genotypes (G), temperatures (T) or significant interaction between G and T (GxT) for a given day using two-way ANOVA (based on the general linear model) corrected with Geisser-Greenhouse at *P* < 0.05. d0, d1, d13 indicate the number of days in DV condition. Letters on the outside of the boxes to the right indicate statistically different groups at *P* < 0.05 as determined by three-way ANOVA followed by a Tukey test. The dashed horizontal line indicates the theoretical limit. (**E**) Maximum net CO_2_ assimilation rate and (**F**) Optimal temperature of net CO_2_ assimilation rate after polynomial fitting of thermal response curves. Asterisks indicate statistically significant differences between genotypes (G), days (D), or significant interaction between G and D (GxD) using two-way ANOVA at *P* < 0.0001 (****). Letters indicate statistically different groups at *P* < 0.05, as determined by two-way ANOVA followed by a Tukey test. For D-F, *n* = 1 leaf per experiment, *N = 6-8* independent experiments.

### Exponential growth of *T. oblongifolia* at 47°C is enabled by photosynthetic heat tolerance

Photosynthesis is one of the most heat sensitive physiological processes in plants (*20, 21*). To test if photosynthetic heat tolerance is responsible for *T. oblongifolia*’s exponential growth in DV summer, we measured leaf-level gas-exchange under standard conditions (day 0), and after 1 and at least 13 days under DV summer (Fig. 1A). When plants were grown under standard conditions, net CO_2_ assimilation rate started declining at ∼38°C (Fig. 1D-F). No significant difference was detected between the photosynthetic performance of *T. oblongifolia* and *A. hypochondriacus* under standard conditions. The net CO_2_ assimilation rates of *A. hypochondriacus* collapsed within a day under DV summer (Fig. S7). No acclimation was observed in *A. hypochondriacus* even after several days under these conditions (Fig. S5). In contrast, in response to DV summer, net CO_2_ assimilation rate of *T. oblongifolia* increased significantly within 13 days (Fig. 1D, Fig. S7-S8, Data S3). The two accessions (DV, DP) behaved similarly in these responses. This is consistent with the hypothesis that *T. oblongifolia*’s exponential growth response to DV summer is enabled by unusual thermal stability of photosynthesis at extremely high temperatures.

To evaluate the speed and mechanisms of photosynthetic acclimation in *T. oblongifolia*, the maximum photosynthetic rate and the temperature at which highest photosynthetic rate was observed (optimal temperature of net rate of CO_2_ assimilation) were evaluated for each accession and condition (Fig. 1E-F, Fig. S9, Data S3). DV summer induced a significant upward shift in the optimal temperature of CO_2_ assimilation rate within a day (Fig. 1F). The highest optimal temperature and a significant increase of CO_2_ assimilation rate were acquired within 13 days. Stomatal conductance did not increase within a day, suggesting that the upward shift in the optimal temperature of CO_2_ assimilation rate observed within a day is controlled by photosynthetic biochemical components rather than by stomatal processes (Fig. S8). After 13 days under DV summer, fluorescence-derived estimates of linear electron transport increased continuously up to 50°C, and the ratio between linear electron transport and net CO_2_ assimilation decreased to nearly the theoretical limit of 4 e^-^ through the linear pathway per CO_2_ fixed (Fig. 1D) (*22*). These patterns indicate that heat acclimation involves stabilizing linear electron transport to a greater degree than downstream processes of photosynthesis and increasing the efficiency with which electron transport is coupled to carbon fixation. How *T. oblongifolia* achieves this remarkable thermal optimum and heat tolerant CO_2_ assimilation is not yet known.

### Photosynthetic heat tolerance of *T. oblongifolia* is greater than that of any other higher plant examined thus far

To compare the photosynthetic heat tolerance of *T. oblongifolia* with that of other higher plants, we compared published thermal optima of net CO_2_ assimilation in 69 plants from the literature with the 2 accessions each of *T. oblongifolia* and *A. hypochondriacus* in this study and expanded previous analyses (*20, 23*). Optimal photosynthetic temperature was significantly correlated with the average maximum temperature of the native habitat across these plants (Fig. S10A, Data S3). We confirmed that thermotolerance limits are variable among plant species and generally reflect the maximum average temperature of their native environment. In addition, *T. oblongifolia* exhibited one of the highest thermal optima of photosynthesis among these plants (Fig. S10B, Data S3). The optimal temperature for photosynthesis acquired after acclimation to DV summer was 45°C (Fig. 1F), which was close to the optimal temperature for photosynthesis measured *in situ* (47°C) (*14*) and to the maximum average temperature of its native environment (46.7°C) (Fig. S1). At 45°C, the maximum CO_2_ assimilation rate of *T. oblongifolia* was higher than that of major crops at more moderate temperatures (Fig. S10C, Data S3). This demonstrates the remarkable photosynthetic rate and thermal stability of *T. oblongifolia* to thrive in conditions currently unsustainable for most plants.

### Photosynthetic acclimation to Death Valley summer is associated with increased density of chloroplasts and mitochondria

Photosynthesis occurs in chloroplasts, which are highly sensitive to heat stress (*24*). To assess the response of chloroplasts to DV summer, we used live imaging to characterize the chloroplast population in mesophyll (M) and bundle sheath (BS) cells surrounding tertiary veins of *T. oblongifolia* leaves challenged with DV summer (Fig. 2A, Fig. S11A, Data S4). We observed an increase in chloroplast density in both cell types in response to DV summer, primarily due to reduced cell volume. Mesophyll cells increased chloroplast density much more than BS cells (Fig. 2A). Interestingly, the leaves that formed after being transferred to DV summer were smaller (Fig. S6B). The reduction in cell size may explain the reduction in leaf size developed under DV summer. Mitochondria generate energy for the cell and contribute to minimizing the effect of disrupted photosynthesis (*25*). Mitochondrial electron transport also participates in the regulation of the cell redox balance during photosynthesis (*25*). To assess the effects of DV summer on mitochondria, we characterized mitochondrial population using rhodamine 123, a mitochondria-specific fluorescent stain (Fig. 2B, Fig. S11B, Data S4). Similar to chloroplasts, DV summer increased the density of mitochondria of M cells in both accessions. Interestingly, in BS cells, mitochondrial density increased only in the acclimated DV accession. Cytoplasm of metabolically active plant cells with high demand for ATP, such as transfer cells and companion cells in the vasculature, contains a high proportion of mitochondria (*26*). This suggests that M cells of both accessions and BS cells of the DV accession may be more metabolically active in DV summer than in standard conditions.

**Fig. 2.**
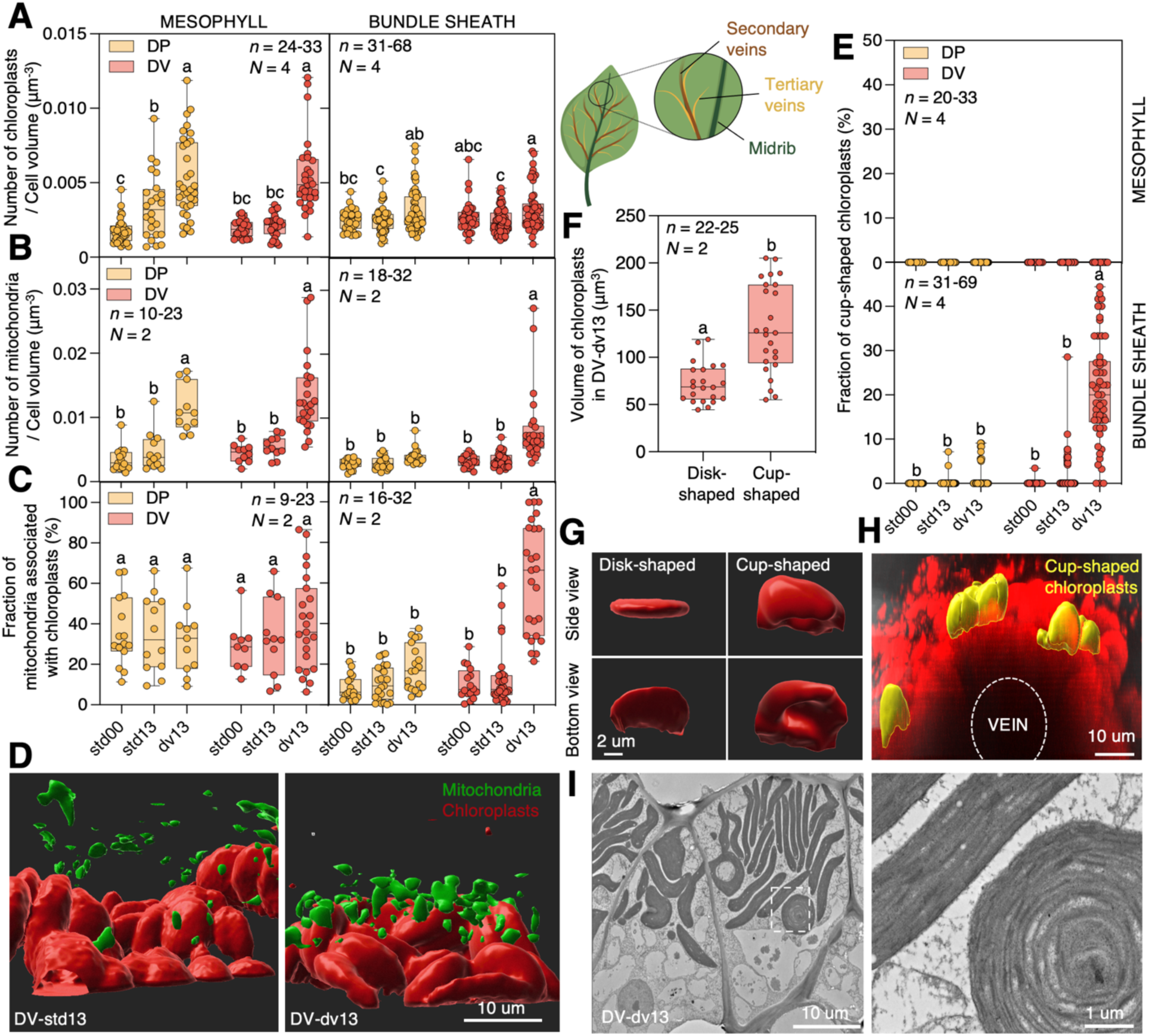
Characterization of chloroplasts and mitochondria in tertiary veins of *T. oblongifolia* challenged with DV summer. All data in this figure are from tertiary veins in DP and DV accessions grown under standard (std13) or DV summer (dv13) for 13 days. std00 indicates the day before plants were moved to the DV condition. A cartoon depicting a venation pattern. *n* = number of biological replicates of *N* independent experiments. **(A)** Quantification of chloroplasts per cell volume in M and BS cells. Letters indicate statistically different groups at a *P* < 0.05, as determined by two-way ANOVA followed by a Tukey test. **(B)** Quantification of mitochondria per cell volume in M and BS cells. **(C)** Number of mitochondria associated with chloroplasts relative to the total of mitochondria per M and BS cell. **(D)** 3D reconstruction using confocal micrographs of leaf BS cells, visualizing chloroplast autofluorescence (red) and mitochondria stained with rhodamine123 dye (green). **(E)** Quantification of the number of cup-shaped chloroplasts relative to the total number of chloroplasts per cell in M and BS cells. **(F)** Volume characterization and **(G)** 3D reconstruction of disk- and cup-shaped chloroplasts presented with a side view and bottom view of acclimated DV accession (dv13). **(H)** Live cell imaging showing orientation of cup-shaped chloroplasts (yellow) in BS cells. **(I)** Representative TEM images of BS cells from DV accessions grown under DV summer for 13 days (dv13) with a magnification on the right.

### The DV accession differs from the DP accession at the subcellular level

To look for any additional differences in the anatomical changes between accessions in response to DV summer, we investigated structural reorganization of chloroplasts and mitochondria of cells near tertiary veins of leaves using live imaging. Only in the acclimated DV accession, organelles reduced volume (Fig. S11) and mitochondria relocated significantly to chloroplasts (Fig. 2CD, Data S4). In addition, we observed an unusual chloroplast with a cup-shaped structure preferentially present in the BS cells of acclimated DV accession (Fig. 2E, Fig. S12, Data S4). While the cup-shaped chloroplasts accounted for ∼20% of all chloroplasts in BS cells near midrib, secondary veins, and tertiary veins of the acclimated DV accession, they accounted for ∼13% in midrib and secondary veins and less than 2% in tertiary veins of the acclimated DP accession (Fig. S12). No cup-shaped chloroplasts were found in M cells (Fig. 2E, Data S4). The volume of cup-shaped chloroplasts was larger than that of the disk-shaped chloroplasts (Fig. 2FG, Data S4) with the cup opening directed primarily toward the vein (Fig. 2H). Cup-shaped chloroplasts showed concentric rings of thylakoid membranes in transmission electron microscopy (Fig. 2I). Overall, these observations showed that while DV and DP accessions harbored similar growth and photosynthesis performance under DV summer, differences could be observed at the subcellular level.

### *Tidestromia oblongifolia* rewires its transcriptome in response to Death Valley summer

We observed physiological, morphological, and anatomical changes in *T. oblongifolia* when challenged with DV summer (Fig. 1-2). To find the genetic determinants of these changes, we sequenced its genome and profiled its transcriptomic changes in response to DV summer. Genomic DNA of the both accessions were sequenced and assembled. Alignment of all contigs from both accessions showed an average pairwise identity of 99.62%. In this study, we used the DP genome for all gene model predictions and expression analyses. The DP genome size is 2.16 Gb with contig N50 of 9.19 Mb and a BUSCO genome completeness score of 97.8% (Tables S1-2). A total of 38,839 protein-coding genes were identified. Gene function annotations of predicted proteins were assigned using Interproscan (*27*), with 37%, 57%, and 48% of the predicted proteins assigned with Gene Ontology, Panther, or Pfam terms, respectively.

To understand how the transcriptome changed during acclimation to DV summer, we collected fully expanded leaves when photosynthetic performance of *T. oblongifolia* was highest on days 0, 1, 5, 13, and 18 after challenging the plants with DV summer (Fig. S13, Data S3), and performed RNA sequencing (RNA-seq) analyses. We identified 9,282 differentially expressed genes (Fig. S14) that were grouped into 8 clusters. 4,162 genes increased their transcript abundance in response to DV summer (clusters 1-4), 3,753 decreased (clusters 5-6), and 1,367 were differentially expressed between the two accessions but not responsive to DV summer (clusters 7-8). Nearly 85% of the genes followed the same pattern between the two accessions (clusters 1-6) and the overall profiles were not significantly distinguishable between 5, 13, and 18 days under DV summer (Fig. 3A-C, Data S5). Most transcriptional changes occurred within a day of the transfer to DV summer, except the genes in cluster 4 that increased their expression more gradually during the first 2 weeks in DV summer. The most immediate increases in transcript abundance (cluster 1) came from genes whose functions were enriched in protein folding, heat response, polyol biosynthesis, and cytidine deamination. More moderately up-regulated expression changes (cluster 2) came from genes whose functions were enriched in response to light stimulus and transmembrane transport. Gradual increases of gene expression were enriched with genes involved in carbohydrate metabolic processes (cluster 4). This is consistent with the growth acclimation responses observed within 10 days (Fig. 1). Down-regulated genes were enriched in catabolic processes, hormone-mediated signaling pathways, and phosphorelay signal transduction system (Fig. 3D, Data S5).

**Fig. 3.**
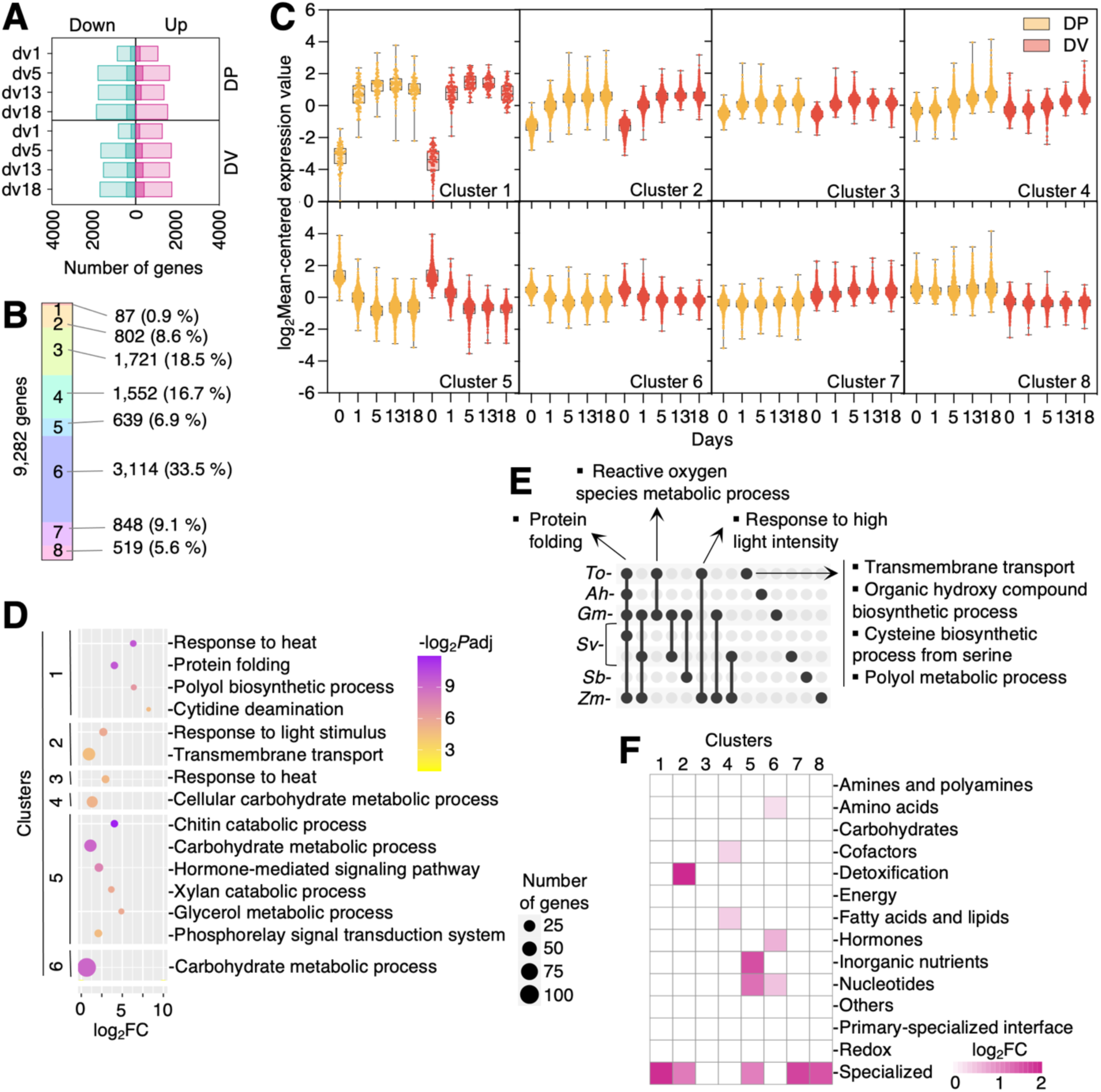
Global gene expression analysis in response to DV summer. **(A)** Global expression analysis of up- and down-regulated genes in response to DV summer for Dos Palmas (DP) and Death Valley (DV) accessions. dv1, dv5, dv13, dv18 indicate the number of days in DV summer. The number of genes displaying at least a two-fold change with false discovery rate (FDR) < 0.01 is shown. Metabolic genes are highlighted as darker green and magenta. **(B)** Number, distribution, and **(C)** Expression profiles of differentially expressed genes grouped by cluster. **(D)** Bubble plots showing significant enrichment of GO biological processes in each cluster (without redundant terms) relative to all expressed genes. Colors and bubble sizes indicate FDR-adjusted –log_2_*P* values and number of genes, respectively. Fisher’s exact tests were performed, and *P* values were corrected for multiple comparisons using the False Discovery Rate (FDR). No enrichment was found in clusters 7 and 8. **(E)** An UpSetR plot showing enrichment in GO biological processes in genes up-regulated in response to high light and high temperature of this study and curated articles. Species are plotted horizontally and intersections of the GO terms between species vertically. GO terms enriched specifically in *T. oblongifolia* are detailed next to arrows. *To*, *Ah*, *Gm*, *Sv*, *Sb*, and *Zm* stand for *Tidestromia oblongifolia*, *Amaranthus hypochondriacus*, *Glycine max*, *Setaria viridis*, *Sorghum bicolor*, and *Zea mays*. **(F)** A heatmap displaying significant enrichment in the metabolic domains of each cluster relative to all expressed metabolic genes.

To determine if *T. oblongifolia* exhibited unique signatures to high light and high temperature in their transcriptomic profiles, we compared GO enrichments of up-regulated genes in response to several days of DV summer to other plants challenged with extreme heat and/or high light conditions. The following processes were enriched specifically in *T. oblongifolia* relative to other plants: transmembrane transport, polyol metabolic processes, organic hydroxy compound biosynthetic process, and cysteine biosynthetic process from serine (Fig. 3E, Data S5). This suggests that *T. oblongifolia* may have optimized mechanisms for maintaining ionic and cellular homeostasis as well as protein and membrane integrity in response to heat. Improving control of the redox state, repair system, and increased heat stability of the membrane may enable the high photosynthetic electron transport rate in acclimated *T. oblongifolia,* which was not affected even when the temperature reached 50°C (Fig. 1).

Since most of the enriched GO terms were related to metabolism, we characterized the metabolic response of *T. oblongifolia* in more detail by constructing a genome-scale metabolic pathway database from the *T. oblongifolia* genome. We used the Plant Metabolic Network pipeline (*28*), producing a computationally-predicted representation of the *T. oblongifolia* metabolism. Using this database (TidestromiaCyc 1.0.0), we performed a metabolic domain enrichment analysis. We found that gene clusters up-regulated within a day under DV summer (clusters 1-3) were enriched in specialized and detoxification metabolic domains. The more gradually up-regulated gene clusters in response to DV summer (cluster 4) were enriched in cofactors and fatty acids/lipids metabolic domains (Fig. 3F, Data S5). Individual metabolic pathways highly up-regulated under DV summer belonged to degradation of aromatic compounds and specialized metabolites, salicylate glycosylation, NAD(P)H repair, and detoxification (Fig. S15, Data S5). These pathways may provide an advantage to *T. oblongifolia* in combating the detrimental effects of high temperature and light on membranes and proteins (*29, 30*) and improving photosynthetic efficiency (*31–33*). Overall, these results showed that the acclimation of *T. oblongifolia* to DV summer was correlated with a transcriptional up-regulation of mechanisms of protection, repair, and detoxification systems within a day and a more progressive transcriptional up-regulation of carbohydrate and lipid metabolism within days of acclimation.

### Genes encoding C_4_ photosynthesis, Photosystem II, and ATP synthase are induced in *T. oblongifolia* in response to Death Valley summer

Acclimation of *T. oblongifolia* to DV summer was correlated with up-regulation of carbohydrate metabolism (Fig. 3) and increased rates of electron transport and CO_2_ assimilation (Fig. 1). To identify potential key determinants of thermotolerance, these pathways were studied in detail (Fig. 4A, Fig. S16, Data S6). Expression of the genes involved in C_4_ photosynthesis was significantly increased in response to DV summer, which may contribute to the high CO_2_ assimilation under DV summer (Fig. 1). Nearly 60% of the nuclear-encoded subunits of Photosystem II and all of the nucleus-encoded subunits of ATP synthase were significantly increased in transcript abundance in response to DV summer. Interestingly, among the Photosystem II subunits, psbS, which plays a major role in photoprotection (*34*), was one of the most up-regulated genes in response to DV summer in our study. Photosystem I is more heat-stable than Photosystem II (*8, 35*). Consistently, Photosystem II was more significantly altered by DV summer than Photosystem I based on the expression of their nucleus-encoded subunit genes. Thus, gene expression regulation of nuclear-encoded subunits appears to be correlated with thermostability of the photosystems.

**Fig. 4.**
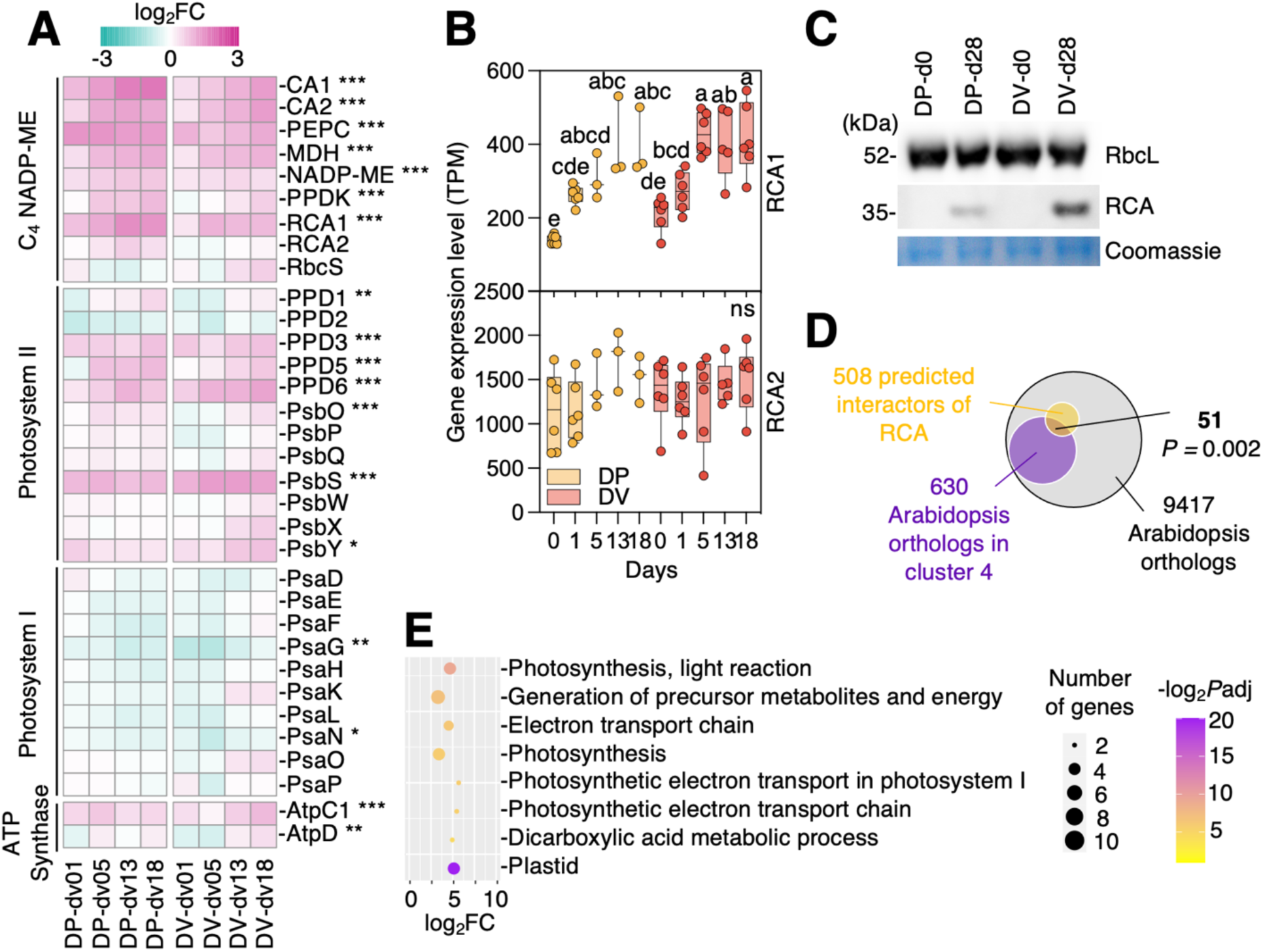
Possible genetic determinants of thermotolerance in *T. oblongifolia.* (**A**) A heatmap displaying the log_2_ fold change in expression of NADP-ME C_4_ and photosynthetic electron chain pathway genes in response to 1, 5, 13, and 18 days (dv01-dv18) of DV summer in DP and DV accessions. Asterisks indicate statistically significant differences between environmental conditions using two-way ANOVA at *P* < 0.05 (*), < 0.01 (**), < 0.001 (***). *P* values were corrected for multiple comparisons using False Discovery Rate (FDR). Full names of proteins encoded by the genes are listed in Data S6. **(B)** Gene expression level expressed in TPM (Transcripts Per kilobase Million) of RCA1 and RCA2 in response to DV summer for DV and DP accessions. Same conventions for labeling as Fig. 2A. (**C**) Leaf proteins of DP and DV accessions grown under standard conditions or under DV summer for 28 days, analyzed by Western blot using RbcL (Rubisco Large unit) and RCA antibodies. (**D**) A Venn diagram showing enrichment of predicted interactors of RCA in the cluster 4. Fisher’s exact tests were performed. (**E**) Significant enrichment of GO biological processes and cellular component in the set of 51 predicted interactors of RCA shown in (D) relative to all expressed orthologs of Arabidopsis.

### Rubisco activase is induced in response to Death Valley summer at the transcript and protein levels

*T. oblongifolia*’s Rubisco was expressed stably in response to DV summer (Fig. 4A). However, in heat-sensitive plants, Rubisco activity decreases under heat stress (*36, 37*). Rubisco activity is regulated by Rubisco activase (RCA) that can activate and repair Rubisco (*38, 39*). To examine the role of RCA on thermotolerance in *T. oblongifolia*, we investigated the transcript and protein level responses of RCA to DV summer. *T. oblongifolia* has two copies of RCA, RCA1 and RCA2, which arose from recent gene duplication (Fig. S17, Data S6). RCA1 had lower expression but was significantly induced by DV summer. RCA2 had higher expression that remained unchanged under DV summer (Fig. 4B). Both genes were predicted to encode proteins of ∼52 kD. However, the RCA protein was detected at ∼35 kDa (Fig. 4C, Fig. S18). Post-translational proteolytic cleavage may account for this molecular weight difference between predicted and observed (*40*). The protein was highly induced in response to DV summer in both accessions (Fig. 4C, Fig. S18). To identify other proteins that may work with RCA in *T. oblongifolia*, we assessed the expression patterns of homologs of predicted RCA interactors in *A. thaliana*. Putative interactors of RCA were enriched in gene expression cluster 4 (51/508 genes, Fisher’s exact test, *P* value = 0.002) (Fig. 4D, Data S6). These 51 genes were enriched in biological processes related to the light reactions of photosynthesis, generation of precursor metabolites and energy, electron transport chain, and dicarboxylic acid metabolic process (Fig. 4E, Data S6). These results suggest that RCA could play an important role in photosynthetic acclimation to DV summer.

## Discussion

Heat stress is a major cause of yield loss in agriculture. Temperatures of above 35°C have detrimental effects on growth, development, and yield of most crops (*6*). Here, we dissected a thermophilic plant, *T. oblongifolia,* that thrives at temperatures above 45°C and shows characteristics of thermoadaptation at the whole organism, organ, cellular, and molecular levels. Understanding how *T. oblongifolia* acclimates and thrives under elevated temperatures may offer new ways for designing heat tolerance in crop plants.

*T. oblongifolia* thrives at temperatures unsustainable for most plants (Fig. 1). We observed an exponential growth rate, accompanied by an increase in the rate of linear electron transport, CO_2_ assimilation, and stomatal conductance in response to Death Valley (DV) summer (Fig. 1). Under DV summer, *T. oblongifolia* increased transcript abundance of detoxification and repair pathways, and synthesis of protective components such as heat shock proteins, osmo-protectants (osmolytes, antioxidants, soluble sugars), and specialized metabolites (Fig. 3). These strategies are not unlike those adopted by many other heat-tolerant plants (*41, 42*). The novelty lies in the fact that *T. oblongifolia* can maintain its performance at temperatures above 45°C. In acclimated *T. oblongifolia*, no damage to the thylakoids was observed (Fig. 2I). In addition, no decline in electron transport rate was observed even when the temperature reached 50°C (Fig. 1) and the maximum CO_2_ assimilation rate of *T. oblongifolia* was higher at 45°C than what other productive crops can achieve at more moderate temperatures (Fig. S10C).

In addition to harboring heat-tolerant traits found in other plants, *T. oblongifolia* exhibits a combination of strategies that may favor photosynthetic performance, which have not been described before to our knowledge. First, chloroplast density increased in mesophyll (M) and bundle sheath (BS) cells, primarily driven by a reduction of cell size. Number and density of organelles in BS cells appear to be reliable anatomical criteria for determining photosynthetic capacity of a plant (*43*), consistent with what we observed in *T. oblongifolia*. While leaf reduction (*18, 19*) and structural reorganization of chloroplasts (*44–48*) in response to heat stress have been documented, little is known about changes in cell size and organellar number and density in terms of thermotolerance mechanisms.

Second, *T. oblongifolia* exhibits unusual cup-shaped chloroplasts in BS cells under DV summer. Cup-shaped chloroplasts are larger than disk-shaped chloroplasts (Fig. 2). Cup-shaped chloroplasts have previously been observed in algae (*49*) and *Selaginella* (*50*) that have single chloroplasts per cell, but multiple cup-shaped chloroplasts in the same cell have not been identified previously. Their specific role in *T. oblongifolia* remains to be determined. Organelle shapes are flexible and can change based on abiotic factors (*51*). Spheroidal shapes have a minimal surface area to volume ratio (*51*). A cup shape may increase the surface area to volume ratio compared to a disk and may therefore enhance the interactive surface between the chloroplast and its surroundings and improve the exchange of metabolites. Specifically, we postulate that the pocket formed by the cup-shaped chloroplast may facilitate recapture of the CO_2_ transported into the bundle sheath by the C_4_ cycle for CO_2_ assimilation. In theory, fixation of one molecule of CO_2_ through the C_4_ pathway requires a minimum of 2 ATP to drive one turnover of the C_4_ cycle and another 2 NADPH and 3 ATP to drive one turnover of the C_3_ cycle. The most efficient way to generate this energy is by driving 4 e^-^ through linear electron flow and another 2 e^-^ through NDH-type cyclic electron flow (*22*). However, by translocating CO_2_ from M to BS cells, the C_4_ cycle produces a large concentration gradient, leading to diffusive leakage of CO_2_ from BS back to M cells. The C_4_ cycle is thus required to operate at a higher rate than the C_3_ cycle to support steady-state CO_2_ assimilation. In the experiments presented here, the efficiency of coupling between the activity of the electron transport system and net CO_2_ assimilation increased from d0 to d13 in both accessions of *Tidestromia*, consistent with reduced BS CO_2_ leakage. This is demonstrated by decreased distance from the theoretical optimum of 4 μmol e^-^ linear electron transport activity per μmol CO_2_ assimilated (Fig. 1D, lower panels). At d13, the DV accession exhibited higher coupling efficiency between electron transport and CO_2_ assimilation activities combined with a greater abundance of cup-shaped chloroplasts in BS cells compared to the DP accession (Fig. 2E). These data support the hypothesis that the cup-shaped chloroplasts in BS cells function to reduce CO_2_ leakage by creating pockets that trap CO_2_ released by plastidic NADP-ME until it can be fixed by Rubisco. Also, we observed a trend toward a higher biomass in the acclimated DV accession than in the acclimated DP accession in response to 28 days under DV summer (Fig. S6A). It is possible that the presence of cup-shaped chloroplasts may be advantageous for DV accession and may be a novel strategy to favor high photosynthetic performance of *T. oblongifolia* under DV summer.

Third, the DV accession’s BS cells had a higher density and reduced volume of mitochondria compared to the DP accession (Fig. 2, Fig. S11). Mitochondrial behavior and morphology may influence energy production but the mechanisms underlying how their subcellular organization influences organismal physiology are poorly understood (*52*). Mitochondrial density varies based on the energy demands of the tissue in mammalian cells (*53*). For example, under increased energetic demand, density of inner mitochondrial membrane, proportion of inter-mitochondrial junctions, and mitochondrial density are predicted to increase (*52*). Consistent with this, mitochondrial density increased and mitochondrial volume decreased under DV summer in BS cells of DV accession (Fig. 2, Fig. S11), suggesting increased energetic demand of heat-acclimated plants.

Fourth, mitochondria relocated to chloroplasts only in the BS cells of the DV accession. In addition to shape changes, mitochondria can move rapidly (*54–56*) and associate with energy-consuming organelles such as nucleus, endoplasmic reticulum, (*53*), and chloroplasts (*54*). Mitochondria become more static once they associate with chloroplasts (*55, 56*). Co-regulation and interdependence between chloroplasts and mitochondria with exchange of metabolites such as ATP, NAD(P)H, and carbon skeleton were shown previously (*57, 58*). Coordination between mitochondria and chloroplasts may help cope with lesions in energy metabolism (*59*), and mitigate the effects of excess reactive oxygen species production (*60*). Finally, subcellular repositioning of organelles may also reduce CO_2_ leakage between M and BS cells (*61, 62*).

Our study also unveiled candidate genes and pathways of thermotolerance. We identified enzymes of the C_4_ cycle, and nuclear-encoded subunits of Photosystem II and ATP synthase with increased transcript abundance under DV summer. With the exception of RCA and Rubisco, the possible role of these candidates in susceptibility or tolerance to heat is not widely documented. Interestingly, rice plants with increased abundance of the nuclear-encoded subunit of chloroplast ATP synthase (AtpD) have higher CO_2_ assimilation and higher electron transport rate under high CO_2_ and high irradiance (*63*). In addition, nuclear-encoded synthesis of the D1 subunit of Photosystem II enhanced survival rates of Arabidopsis, tobacco, and rice under heat stress (*64*). Characterizing *T. oblongifolia* photosynthetic genes whose expressions were induced may reveal novel mechanisms or targets of thermotolerance.

RCA has been considered to be a target for engineering heat tolerance (*65*). Several studies proposed that RCA may limit photosynthesis under heat stress (*17, 66, 67*). In addition, Rubisco’s activation state constrained the increase in CO_2_ assimilation observed in maize expressing more Rubisco (*67*). Nonetheless, it is still difficult to determine if RCA is the ultimate site of high temperature sensitivity or if it is an intermediate component of a limitation that is ultimately traced to the electron transport system (*68*). Our study showed that gene expression and protein accumulation of Rubisco were not significantly changed in response to DV summer (Fig 4A,C). In contrast, one of *T. oblongifolia*’s RCA-encoding genes, RCA1 (TIOBL00G270050), was increased at the transcript level under DV summer and the RCA protein was also induced (Fig 4A-C). The increase in expression of RCA relative to Rubisco suggests that increased RCA activity is required to maintain a proper activation state of Rubisco under DV summer.

The role of each RCA isoform remains to be determined in *T. oblongifolia*. Most plants express two isoforms of RCA (α and β) that are produced either by alternative splicing of a single gene or encoded from two genes (*65*). The α-isoform is generally larger (45-48 kDa) than the β-isoform (41-43 kDa) and contains a C-terminal domain with two redox-sensitive cysteine residues (*69*). While both copies of *T. oblongifolia* include a C-terminal domain homologous to the *A. thaliana* α-isoform, one of the redox-sensitive cysteines was replaced with tyrosine in RCA2 (TIOBL00G358250) (Fig. S17). RCA1 could therefore be analogous to the α-isoform and RCA2 to the β-isoform due to lack of this redox-sensing residue. Further supporting this hypothesis, previous studies on *S. bicolor* and *S. viridis* found that gene expression of RCA-α was induced by heat treatment (*69, 70*) and basal expression of RCA-β in *S. viridis* was higher than that of RCA-α. Consistently, high temperature also induced the protein level of RCA-α that was undetectable under control conditions in grasses (*69*). The specific role of each isoform in plants remains unclear and may be species-dependent (*65*).

While the interaction of RCA with Rubisco has been well documented (*71*), RCA’s interaction with the thylakoid membrane and its potential role in other processes is poorly understood. RCA has been proposed to be associated with thylakoid-bound ribosomes to protect the thylakoid-associated translation machinery against heat inactivation (*72*). In addition, a recent proteomic study indicated that RCA may be in close association with Photosystem I (*73*). Overexpression of RCA in rice disturbed Photosystem I electron transport (*74*). Other studies suggested a role of RCA in protecting the thylakoid membrane from oxidative stress (*65*). Our study showed that RCA may be a hub of a network that influences various aspects of photosynthesis. Many of its predicted interactors that were up-regulated in response to DV summer are involved in the photosynthetic electron transport chain (Fig. 4). Further characterization of the up-regulated network around RCA may reveal insight into how RCA functions in thermotolerance of photosynthesis.

In conclusion, current predictions of climatic conditions require the development of strategies to improve crop resilience to heat stress. Our holistic dissection of the thermophile *T. oblongifolia* at the whole organism, organ, cell, and molecular levels adds new dimensions to our understanding of thermotolerance. With its genome now sequenced and annotated, *T. oblongifolia* may become a useful model for identifying, selecting, and transferring thermotolerance traits into crop plants.

## Acknowledgments

We would like to thank Drs. Joseph Berry, Harold Mooney, and members of the Rhee lab for their helpful discussions on the project, summer interns Galyna Vakulenko, Bharti Parihar, and Suzie Lee for preliminary experiments on the project, William Dwyer for curation of sorghum C_4_ photosynthesis pathway genes, Michael Pidgeon for preliminary TEM analysis, Stanford Statistics consulting for providing statistical advice, Carnegie facilities managers Theo van de Sande, Giancarlo Materassi-Shultz, and Ismael Villa for general facilities and equipment support, IT managers Garret Huntress and Maria Lopez for IT support, Microscopy manager Andrey Malkovskiy for microscopy assistance, additional seed collectors (Soon Sup Rhee, Colin Jin Roth, Joo Hyun Rhee, Chris Roth, Carol Fields), Death Valley National Park for providing seed collection permits, California Botanic Garden for providing *T. oblongifolia* seeds (accession number: 24132), and National Plant Germplasm System North Central Regional Plant Introduction Station (NPGS NCRPIS, USDA) for providing *A. hypochondriacus* seeds. This work was done on the ancestral land of the Muwekma Ohlone Tribe, which was and continues to be of great importance to the Ohlone people.

## Funding

this work was supported by:

Carnegie Venture Grant (10908) (S.Y.R., J.B., and J.E.J.)

US National Science Foundation grants (MCB-1617020, IOS-1546838) (S.Y.R.)

Water and Life Interface Institute (WALII) DBI (grant no. 2213983) (S.Y.R.)

US Department of Energy, Office of Science, Office of Biological and Environmental Research, Genomic Science Program grant nos (DE-SC0018277, DE-SC0008769, DE-SC0020366, DE-SC0023160, and DE-SC0021286) (S.Y.R.)

The Guangdong Science and Technology Foundation, “ZhuJiang Talent Innovation” project (2019ZT08N628) (S.C.)

The National Natural Science Foundation of China “Excellent Young Talent” (32022006) (S.C.)

GuangDong Basic and Applied Basic Research Foundation (2022A1515110358) (H.L.)

## Author contributions

K.P., J.E.J., S.Y.R. conceived the project; K.P. prepared samples, performed most of the experiments, analyses, data curation, and visualization; K.P. and J.E.J. designed the plant growth conditions and physiological experiments, H.L. and S.C. sequenced and created assembly of genome, B.X. and M.S created annotations of genome and processed RNA sequencing reads, B.X. performed differential expression analysis, K-means clustering, and developed web servers, M.S. conducted the phylogenetic and sequence analyses of RCA genes, S.F. performed live microscopy and image analysis, R.-C.H. performed TEM, C.L.H. created PGDB, K.P. wrote the original draft, S.Y.R., J.E.J., S.F., M.S., C.L.H. edited the manuscript, the manuscript was read by all the authors. S.Y.R. supervised K.P., B.X., M.S., S.F., and C.L.H. S.Y.R., J.E.J., S.C., H.L. acquired funding.

## Competing interests

authors have declared no competing interests.

## Data and materials availability

Data are available in the supplementary materials. FASTQ files of the RNAseq dataset were deposited at NCBI’s Sequence Read Archive (PRJNA970737). Expressions, sequences, and annotations server is available at: https://tidestromia-expression.rheelab.org/ Blast viewer server is available at: https://tidestromia-blast.rheelab.org/ TidestromiaCyc 1.0.0 is part of the Plant Metabolic Network and is available at: https://pmn.plantcyc.org/organism-summary?object=TIDESTROMIA

Bespoke scripts were deposited at: https://github.com/TheRheeLab/Tidestromia

## Notes

https://tidestromia-expression.rheelab.org/

https://tidestromia-blast.rheelab.org/

https://pmn.plantcyc.org/organism-summary?object=TIDESTROMIA

https://github.com/TheRheeLab/Tidestromia

### Competing Interest Statement

The authors have declared no competing interest.

### Summary of Updates

Author affiliations updated; Materials and methods updated; Supplemental files updated; Acknowledgements updated

