## Supplementary_materials for "Photosynthetic acclimation mediates exponential growth of a desert plant in Death Valley summer"

**The PDF file includes:**

Materials and Methods
Figs. S1 to S18
Tables S1 to S3

**Other Supplementary Materials for this manuscript include the following:**

Data S1 to S6

### Materials and Methods

Seed collection and permits

Seed collection of *T. oblongifolia* at Death Valley was authorized by the United States Department of the Interior National Park Service (Death Valley) under the permits DEVA-2018-SCI-0047 and DEVA-2018-SCI-0055 for the study DEVA-00522. A maximum of 1,000 seeds were collected close to Furnace Creek (CA) (36.403211, -116.777922) and no more than 10% of the seeds from a single plant was collected. In the beginning of the project, we obtained seeds from California Botanic Garden (CA) collected near the Dos Palmas area (33.55202, -115.80580). Additional seed collection of *T. oblongifolia* at Dos Palmas was done outside the area of a National Park (33.322051, -115.996401) (Fig. S4). We followed the same rules as those described in the Death Valley permit for all our collections. The permit reports are available on the National Park Service Research Permit and Reporting System (RPRS) portal. Seeds of *Amaranthus hypochondriacus* (Nebraska accession PI558499, India accession PI480569) were obtained from the National Plant Germplasm System (NPGS, USDA).

Plant growth

***Standard growth conditions.*** *Tidestromia oblongifolia* seeds were germinated in controlled environmental standard conditions (50%HR, 16h/8h day/night with 400 μmol $m^{-2}$ $s^{-1}$, and 31ºC/22ºC day/night temperature, dawn 6 am, dusk 10 pm) in small pots (6.0 x 8.5 x 5.5 cm, T.O. Plastics 715399C-BX, Clearwater, MN, USA) filled with dry soil mixture with a ratio of: 2 scoops (3-quart scoop) soil (Promix HP 20381, Premier Horticulture, Quakertown, PA, USA), 1 scoop turface (Turface Athletics MVP, Profile Products LLC, Buffalo Grove, IL, USA), and 1 small teaspoon (15 mL) of 14:14:14 osmocote slow-release fertilizer (Osmocote 14-14-14 F1877, Everris NA, Dublin, OH, USA). Pots were covered with a clear humidity dome (CN-DOME, Greenhouse Megastore, West Sacramento, CA, USA) and seedlings were irrigated with a few drops of water every two days. Around 3 weeks after germination, seedlings were transplanted into deeper pots (12.7 x 12.7 x 30.5 cm, Stuewe and Sons 1-85682, Tangent, OR, USA) with the same ratio of dry soil mixture. Seedlings were irrigated with water before and after the transfer by placing the pots inside 25.4 x 53.4 x 5.1 cm trays (CN-FLXHD-X2, Greenhouse Megastore, West Sacramento, CA, USA) and filling the trays with water. Around 6 weeks later*,* the plants either remained in the standard conditions or were transferred into a separate, controlled environmental chamber that mimicked Death Valley conditions (described below). Plants growing in deeper pots were watered frequently to ensure that water was always present in the bottom trays. *Amaranthus hypochondriacus* seeds were germinated in the same conditions as *T. oblongifolia* seeds. Around 2 weeks after germination, seedlings were transplanted into deeper pots with the same ratio of soil mixture. Around 4 weeks later*,* the plants were transferred to Death Valley conditions the last hour before the transition from day to night to obtain overnight dark acclimated leaves prior to the measurements. As the developmental rate of *A. hypochondriacus* was faster than that of *T. oblongifolia*, plants of different ages were used for *T. oblongifolia* and *A. hypochondriacus* to match the developmental stage of the two species.

***Death Valley (DV) summer growth conditions.*** The Death Valley growth conditions were designed to mimic the solar radiation and air temperature on a typical day in July at Furnace Creek in Death Valley (36.42041°N, 116.84714°W). To estimate the daily cycle of photosynthetically active radiation, total shortwave radiation was modeled following (*75*). To estimate the daily cycle of air temperature, measurements of daily maximum and minimum temperatures from the Furnace Creek, CA weather station were obtained from the Global

Historical Climatology Network (https://www.ncdc.noaa.gov/cdo-web/search?datasetid=GHCND). These were interpolated with an empirical function from (*75*). The radiation and temperature cycles for a typical July and January day were then programmed into growth cabinets (PtGC1-1200H LED Z3WC, HiPointUS and GC2-1200H LED Z4, Taiwan Hipoint Corporation, Kaohsiung, Taiwan), respectively. The daylight (14h/10h day/night) was shifted by 3 hours (from 6 am in the field to 9 am in the custom-built growth chambers). The modeled climate was sufficiently accurate to replicate the light intensity and temperature measurements made in the field in the 1970s (Fig. S1).

Determination of growth rate, biomass, and leaf number

Pots were weighed daily for one month with a scale (Ohaus, model CL500, capacity 5000g x1g, USA) to determine the growth rate. On days 0, 14, and 28 after the transfer to the DV chamber, 2-4 plants were sacrificed for each accession, each condition, and each harvest day. Shoot and root of each plant were separated, weighed to get the fresh weight (FW), then dried in a convection oven at 45ºC until the change in the biomass was less than 1% per hour (around 5 days). All leaves of the plants were collected and leaf area was determined with ImageJ (*76*) after covering the leaf with a glass pane to limit leaf curvature.

Characterization of leaf physiology with chlorophyll fluorescence and gas-exchange

To characterize the physiological performance of leaves, we used a LI-6800 portable gas-exchange system equipped with a pulse-amplitude modulated (PAM) fluorometer 6800-01A (LI-COR Biosciences, Lincoln, NE, USA).

***(1) Adaptation of leaf cuvette.*** A standard leaf cuvette was used with an aperture diameter of 2 or 6 $cm^2$ as appropriate for the surface area of *Tidestromia oblongifolia* and *Amaranthus hypochondriacus* leaves. To catch broken trichomes and keep them out of the flow path, fine mesh gauze was fastened across each aperture and sealed with Apiezon Q putty (M&I Materials, United Kingdom). Before beginning the experiment, blank measurements were performed to verify that the modified apertures provided a leak-free seal.

***(2) Installation of leaf cuvette in growth chamber.*** During the experiment, both the plants and the measuring head of the LI-6800 system were enclosed inside a growth chamber where the environmental conditions could be controlled. This was useful for all the measurements because it stabilized the plant's environment, and it was especially important for the temperature response measurements because it extended the temperature control range of the cuvette.

***(3) Cuvette configuration for measurements.*** For all of the measurements, the actinic light in the cuvette was provided as a mixture of red (90%) and blue (10%) wavelengths (R90B10). The PAM analysis used a multi-phase flash (i.e., three 300 ms phases; first and third phase at 5,000 µmol PPFD $m^2$ $s^{-1}$; second phase ramped down by 25% for determination of Fm′), followed by 2 seconds of far-red illumination and a 5 seconds of dark pulse for determination of Fo′. The cuvette was stirred with fan speed 10,000 RPM and maintained at a flow rate 600 µmol $s^{-1}$. All measurements were performed at 21% oxygen and 25% relative humidity inside the cuvette.

***(4) Photosynthetic response to all factors in the growth environment.*** To determine the daily maximal photosynthetic rate under the growth conditions, we measured single leaves throughout the day while matching the light intensity and air temperature in the cuvette to those in the growth chamber. Plants were transferred from standard into DV summer conditions the last hour before the transition from day to night of DV summer to obtain overnight dark acclimated leaves prior to the measurements. Measurements started the next day within one hour before the transition from night to day. These measurements were performed on 6 leaves from 3 plants/accession/condition. One experiment was conducted in this study for each accession.

***(5) Photosynthetic response to single environmental factors.*** Each plant was transferred into DV summer for 24 hours (day 1) or at least 13 days (day 13) prior to being transferred into a cabinet set at 31ºC for the measurement. Before each measurement sequence, the leaf was permitted to equilibrate to the cuvette environment for 15-30 minutes; and within each sequence, measurements were made at 5 min intervals. To assess the steady-state temperature response, the light was held at 2200 µmol PPFD $m^{-2}$ $s^{-1}$ and the cuvette carbon dioxide at 400 µmol $CO_2$ $mol^{-1}$. The leaf temperature was progressively increased from 31ºC to 50ºC in one-degree steps. At the end of the day of measurements, the leaf was removed from the cuvette and a photograph was taken to determine leaf area within the cuvette. The measurements were performed on 1 leaf per experiment. Six to eight independent experiments were conducted for this study for each accession and condition.

***(6) Determination of maximum net $CO_2$ assimilation rate and optimal temperature of net $CO_2$ assimilation rate.*** For each accession and condition, temperature response curves of net $CO_2$ assimilation rate were fitted with a polynomial equation to determine the maximal net $CO_2$ assimilation rate and the temperature at which highest photosynthetic rate was observed. This temperature refers to the optimal temperature of net $CO_2$ assimilation rate or thermal optimum of photosynthesis.

***(7) Determination of leaf absorptance to photosynthetically active radiation.*** The leaf absorptance was measured between 400-700 nm with an integrating sphere (Analytical Spectral Devices, Inc., Boulder, CO, USA) and spectrometer (AvaSpec-ULS3648, Avantes, Lafayette, CO, USA). To determine the absorptance to the light regime used in the LI-6800 experiments, we used a weighted average with 600-650 nm representing absorptance to the 625 nm LEDs (red, 90%) and 450-500 nm representing the 475 nm LEDs (blue, 10%). These absorptance values were used to estimate the rates of linear electron flow from the fluorescence measurements by assuming that the absorbed light was partitioned evenly between Photosystems I and II.

Comparison of optimal temperatures of photosynthesis

Temperature response curves of net photosynthesis, photosynthetic rate, $CO_2$ uptake, or $CO_2$ assimilation were searched in the literature and optimal temperature of photosynthesis recalculated using ImageJ. The studies (Data S3) were selected on the basis of a detailed description of the location (region and country) of measurements in the field. Based on the location described, a NOAA station close to the site of measurements was searched and the average maximal temperature from 1948 to 2022 was calculated. If a nearby NOAA station was not found, we searched the temperature online using the Google search engine with the names of the region and country of sampling and used the results from the NOAA database obtained from the Google search. Plant categories were attributed in accordance with (*15*, *77*).

Genome sequencing and annotation

***Sample collection.*** Samples were collected from young leaves of the DP and DV accessions grown in standard conditions for 9 weeks and treated with 30h of darkness to reduce the quantity of starch and polysaccharides that may interfere with high-quality genomic DNA extraction. Fresh and flash frozen tissues were shipped to BGI Qingdao (China) for genome sequencing.

***Genome sequencing.*** High-quality genomic DNA was extracted from leaves using the cetyltrimethylammonium bromide (CTAB) method and used to construct a DNA Nanoball (DNB) library for short read sequencing, and a PacBio library for single-molecule real-time (SMRT) sequencing. For DNB-seq, the DNA was fragmented to 300-500 bp in size, amplified to a nanoball, sequenced on DNB-seq platform with the Paired End 150 strategy, which generated 155.41 Gb of sequence. For the PacBio library, the CLR type library was constructed according to the PacBio manual, which generated 158.34 Gb of sequence (Tables S1-2).

***Genome size estimation.*** Short-read data was estimated by gce (v1.0.0) with the following parameters: -k 21 -a 0 -d 0 for kmer_freq_hash and -m 1 -b 1 for gce (*78*) (Table S3).

***Genome assembly.*** The PacBio reads of CLR type were assembled by Canu (v2.0) with the following parameters: genomeSize = 2.17g, minOverlapLength = 700, minReadLength = 1000 (*79*). The bubble sequences of the assembled contigs were removed by minimap2 (v2.1) with parameters -x asm5 and purge_dups (v1.2.3) with -T 2. Then, the contigs were polished with short-read data for two rounds to correct the erroneous SNPs and Indels using Pilon (v1.24) (Table S3).

***Genome repeat identification.*** The EDTA (v1.9.6) and RepeatMasker (v4.1.2) software were used to identify repetitive sequences in the genome of *T. oblongifolia (80)*. Pseudomolecule was first trained by EDTA to generate each library file of transposon elements with parameters --anno 1 --force 1 --debug 1 --sensitive 1 --evaluate 1. Then library files were used as input for Repeatmasker with parameters -a -html -gff (Table S3).

***Gene model prediction.*** The Maker pipeline (*81*) was used to annotate protein-coding genes. The pipeline integrates transcriptome evidence, protein evidence, and *ab initio* prediction. For transcriptome evidence, the transcripts were *de novo* assembled from mRNA-seq reads by HISAT2 (*82*), Trinity (*83*), Cufflinks (*84*), StringTie (*85*), Mikado (*86*), and PASA (*87*) software. Protein evidence includes the predicted proteins generated with Fgenesh (*88*) and homologous proteins downloaded from public data. *Ab initio* prediction was conducted with mRNA-seq data using Augustus, GeneMark-ET, and Braker2 (*89*) (Table S3). Gene IDs followed the nomenclature TIOBL00G000000 (TI = genus, OBL = species, 00 = chromosome, 00000 = number). Because the current release is not yet chromosome-scale, all chromosome numbers were set to 00.

***Genome annotations.*** Gene function annotations of predicted peptide sequences were performed using Interproscan (*27*) with the default behavior of using all available annotation methods (Table S3).

RNA extraction

Fully developed leaves were collected on days 0, 1, 5, 13, and 18 at 6 hours after the night/day transition (zeitbeger time 6) for plants under standard condition and 4 hours after the night/day transition (zeitbeger time 4) for plants under DV summer condition. These time points to collect samples for transcriptomics were determined by comparing diurnal photosynthetic performances of *T. oblongifolia* accessions to *A. hypochondriacus* accessions challenged with Death Valley conditions. The time of peak of performance was identified by assaying photosynthesis rates throughout the day and was selected for sample collection. Tissues were frozen immediately in liquid nitrogen. Samples of *A. hypochondriacus* were not collected beyond 5 days because the plants were dying. Total RNA was extracted using the RNeasy Plant Mini Kit (Qiagen, Valencia, CA, USA) with on-column DNase (Qiagen, Valencia, CA, USA) digestion as described by the manufacturer.

cDNA library preparation and high-throughput sequencing

Quality of RNA extraction was checked by bioanalyzer (RI>8) and samples were sent to Novogene (Sacramento, CA, USA) for QC check and sequencing. All samples passed through the following three steps before library construction: 1) preliminary quantitation by Nanodrop (Thermofisher, Waltham, MA, USA), 2) tests of RNA degradation and potential contamination by agarose gel electrophoresis, and 3) check of RNA integrity and quantification by 2100 Bioanalyzer (Agilent, Santa Clara, CA, USA). After the QC procedures, mRNA was enriched using oligo(dT) beads. The mRNA was fragmented randomly by adding a fragmentation buffer, then the cDNA was synthesized by using mRNA template and random hexamer primers. After first-strand synthesis, a custom second-strand synthesis buffer (Illumina, San Diego, CA, USA) was added, with dNTPs, RNase H, and DNA polymerase I to generate the second strand by nick-translation. AMPure XP beads (Beckman Coulter, Beverly, USA) were used to purify the cDNA. The final cDNA library was ready after a round of purification, terminal repair, A-tailing, ligation of sequencing adapters, size selection, and PCR enrichment. Quality control of the library consisted of testing the library concentration preliminarily by Qubit 2.0 (Thermofisher, Waltham, MA, USA), testing the insert size by 2100 Bioanalyzer, and quantifying the library effective concentration precisely by qPCR. The qualified libraries were fed into Illumina Novaseq 6000 sequencers after pooling based on its effective concentration and expected data volume. Distribution of sequencing quality, error rate, and GC content distribution were checked. The sequenced reads (raw reads) often contain low quality reads and adapters, which will affect analysis quality. Raw reads were filtered to get the clean reads. The filtering process was as follows: 1) remove reads containing adapters, 2) remove reads containing N > 10% (N represents a base that cannot be determined), and 3) remove reads for which low-quality (Qscore<= 5) bases comprise more than 50% of the total bases. The original raw data from the Illumina platform were transformed to Sequenced Reads, known as Raw Data or Raw Reads, by base calling. Raw Data were recorded in a FASTQ file, which contains sequencing reads and corresponding sequencing quality.

Processing of RNA sequencing reads

Highly similar sequences in the genome annotations were removed using CD-HIT (*90*) v4.8.1 with the parameters -c 0.99 (99% similarity cutoff) -G 0 (local rather than global alignment) and -aS 0.75 (the shorter sequence must align to at least 75% of the longer sequence). This non-redundant genome annotation was used for read mapping via HISAT2 version 2.2.1 (*82*) with scoring parameters --score-min L,0,-0.875, with output piped directly into the samtools version 1.8 (*91*) view command to compress output to BAM format. The Transcripts Per Million (TPM) gene expression metric was used because it eliminates statistical biases inherent in the Reads Per Kilobase Million (RPKM) measure (*78*). Normalization between samples has been shown to be essential in RNA-seq data analysis (*92, 93*). We applied the Trimmed Mean of M-values (TMM) normalization method according to (*92*) in order to normalize the expression of genes across samples. TMM generates a normalization factor based on the assumption that most genes should not be differentially expressed between samples (*92*) (Fig. S14).

Pathway Genome Database (PGDB) construction for *T. oblongifolia*, TidestromiaCyc

The PGDB for *Tidestromia oblongifolia* was generated by means of a three-stage pipeline. In the first stage, protein sequences from the *T. oblongifolia* genome were used as input for the Ensemble Enzyme Prediction Pipeline (E2P2) software v4.0 (*94*). This software predicts enzyme functions based on weighted results of BLAST and PRIAM from the reference protein sequence dataset (RPSD) v4.2. RPSD is a database of experimentally-characterized enzyme sequences downloaded from PlantCyc (*94*), MetaCyc (*95*), SwissProt (*96*), and BRENDA (*97*). In the second stage, the predicted enzymes and their associated reactions were used as input for the PathoLogic software, part of the Pathway Tools v24.0 suite from SRI International (*98*). PathoLogic was used to call presence or absence of MetaCyc and PlantCyc metabolic pathways in *T. oblongifolia*, and generate the initial version of TidestromiaCyc based on those calls. In the third stage, TidestromiaCyc was curated using the Semi Automated Validation Infrastructure (SAVI) software v3.1 (*94*), which is used to apply previous manual curation decisions for the appropriate phylogenetic range of each pathway to derive the final PGDB. Finally, experimentally supported pathways and enzymes from MetaCyc and PlantCyc that were curated to be present in *T. oblongifolia* were annotated with the appropriate evidence codes. Transcriptomic data were visualized by using the Omics Dashboard feature of the PGDB provided by Pathway Tools and overlaid onto the pathways and reactions associated with significantly differentially expressed genes. The results were used to generate heatmaps using pheatmap package version 1.0.12 in R.

Gene expression analyses

***Differential gene expression analysis.*** The DESeq2 bioconductor package (*99*) was used to analyze differential expression based on a negative binomial distribution model. The following filtering criteria were used in our analysis: False discovery rate (FDR) < 0.01, $|\log_2(\text{Fold Change})| > 1$, and at least 2 replicates have TPM value > 1. Expression values were $\log_2$-transformed and mean-centered for normalization.

***K-means clustering.*** Genes within the same cluster exhibit the same trends in expression levels. To determine the optimal number of clusters for K-means clustering, we used three different methods, elbow method (*100*), average silhouette method (*101*), gap statistic method (*102*), and the best "K" was selected by visualizing the clusters in 3-dimensional principal component analysis (PCA) space (Fig. S14).

***Enrichment.*** Gene Ontology (GO) enrichment for GO terms under "biological process" (BP), "cellular component" (CC), and "molecular function" (MF) against the expressed genes was conducted using Cytoscape version 3.9.1 with its plugin Bingo version 3.0.3 (*103*). We used GO annotations and ontologies that were released in October 2022. GO annotations were generated from the union of GO terms from Interproscan (version 5.59-91.0 with default parameters and -goterms argument) and GO annotations of *A. thaliana* reciprocal best hit orthologs. To compare GO enrichment of high temperature or high light response genes across plant species, genomes of *Amaranthus hypochondriacus*, *Glycine max*, *Setaria viridis*, *Sorghum bicolor*, and *Zea mays* were downloaded from Phytozome v.13 (*104–109*) and GO enrichment analysis was performed using the up-regulated genes reported in (*70*, *110–112*). For all enrichments, a hypergeometric test was used with a Benjamini and Hochberg FDR correction at the significance level of 0.05 to determine significantly enriched GO terms. Because the transcriptomics were enriched for nuclear-encoded genes, GO enrichment analysis used only nuclear-encoded genes.

***Hand curation of $C_4$ photosynthesis, RCA, Rubisco, and photosynthetic electron chain pathway genes***. $C_4$ photosynthesis and Rubisco genes were manually identified from *T. oblongifolia* and *A. thaliana* by BLASTP (e-value of at least ~1e-5) using the list of $C_4$ genes in *Sorghum bicolor* described in (*113*) as queries. RCA genes were identified from *T. oblongifolia* by BLASTP (e-value of at least ~1e-5) using *A. thaliana* RCA gene (AT2G39730) from TAIR (*114*) as a query. Photosynthetic electron chain pathway genes were manually identified using best reciprocal hits using BLASTP (e-value of at least ~1e-5) between *A. thaliana* and *T. oblongifolia* based on *A. thaliana* genes AT4G15510 (PPD1), AT2G28605 (PPD2), AT1G76450 (PPD3), AT5G11450 (PPD5), AT3G56650 (PPD6), AT5G66570 (PsbO), AT1G06680 (PsbP), AT4G21280 (PsbQ), AT1G44575 (PsbS), AT2G30570 (PsbW), AT2G06520 (PsbX), AT1G67740 (PSbY), AT1G03130 (PsaD), AT2G20260 (PsaE), AT1G31330 (PsaF), AT1G55670 (PsaG), AT1G52230 (PsaH), AT1G30380 (PsaK), AT4G12800 (PsaL), AT5G64040 (PsaN), AT1G08380 (PsaO), AT2G46820 (PsaP), AT4G04640 (AtpC1), AT4G09650 (AtpD) from TAIR as queries. From the matching sequences including $C_4$ photosynthesis in *T. oblongifolia*, we used the genes with the highest expression in all the conditions because $C_4$ isoform genes are highly expressed compared to $C_3$ isoform genes (*115*).

<u>Protein isolation and Western blot analysis</u>

Leaf protein isolation was adapted from (*116*). Briefly, 100 mg of fully developed leaves of plants grown under standard or DV summer conditions for 28 days were collected, frozen immediately in liquid $N_2$, and ground with mortar and pestle. The powder was homogenized in 1 mL of protein extraction buffer (100 mM trisaminomethane-HCl, pH 7.8, supplemented with 25 mM NaCl, 20 mM ethylenediaminetetraacetic acid, 2% sodium dodecyl sulfate (w/v), 10 mM dithiothreitol, and 1⁄2 tablet of protease inhibitor cocktail for 10 mL of solution (Sigma, St Louis, MO, USA). Protein extracts were vortexed for 20 seconds, incubated at 65°C for 10 minutes to denature, vortexed for 20 seconds, and centrifuged at 13,000 g for 1 minute at 4°C to obtain a clear supernatant. Protein extract concentration was evaluated with plate reader Infinite® M1000 (Tecan, Männedorf, Switzerland) using Pierce 660 nm protein assay reagent (22660, Thermofisher, Waltham, MA, USA) supplemented with ionic detergent compatibility reagent (22663, Thermofisher, Waltham, MA, USA). Protein extracts (1 µg or 5 µg) were supplemented with 4x Laemmli sample buffer (1610747, Bio-Rad, Hercules, CA, USA), separated by 12% SDS-PAGE, and transferred onto polyvinylidene fluoride membranes for 2 hours at 75V at 4ºC in transfer buffer (Tris 25 mM, Glycine 192 mM, SDS 0.04%, MeOH 20%). Following transfer, the membranes were washed with Tris-Buffered Saline containing Tween (TTBS; Tris 25mM pH7, NaCl 0.5M, Tween-20 0.1% [v/v]) and saturated with TTBS supplemented with 5% (w/v) skimmed milk for 1 hour at room temperature. The membranes were washed three times, for 20 minutes each, with TTBS and incubated overnight with TTBS supplemented with 1 mg/mL BSA and rabbit primary antibodies at 4ºC. Rabbit primary antibodies against RuBisCO large subunit antibody (Form I) (RbcL, 1:5000, catalog number AS03 037) and anti-Rubisco activase, chloroplastic antibody (RCA, 1:2000, catalog number PHY2139S) were purchased from Agrisera (Vännäs, Sweden) and PhytoAb (San Jose, CA, USA), respectively. The membranes were washed three times for 20 minutes (each time) with TTBS, then rinsed with TBS. Immunodetection was performed using the chemiluminescent protein gel blotting analysis system SuperSignal West Dura (34076, Thermofisher, Waltham, MA, USA) and FluorChemQ gel documentation system for luminescence/fluorescence gel imaging (Alpha Innotec,

Kasendorf, Germany). Unless stated otherwise, chemicals were purchased at Millipore Sigma (Burligton, MA, USA).

Live cell imaging and quantification of organelles

***Leaf dissection.*** Leaf disks (8 mm diameter) were collected using a leather hollow hole punch from freshly harvested leaves. Immediately before imaging, the epidermis and trichomes on both sides of the leaf disk were removed along with the midrib and secondary veins, which was required to allow penetration of the confocal laser necessary for live cell imaging. Following the leaf disk dissection, the leaf disks were briefly washed (~5 seconds in 10 mM MES NaOH pH 5.7 and 10 mM $MgCl_2$) to remove cellular debris.

***Live cell imaging.*** Live imaging was performed using a Leica SP8 scanning laser confocal microscope (Leica, Wetzlar, Germany). Leaf disks were imaged in the X, Y (~200 μm by ~200 μm), and Z (20- to 30- 1 μm Z-sections) planes using a 63x water objective and Hybrid SMD2 detector, with a 1.2 aperture. Chloroplasts were visualized by exciting chlorophyll autofluorescence with a white light laser (506 nm), and fluorescence was detected between 656 nm - 691 nm. Mitochondria were visualized by first staining leaf disks in 0.10 mg/mL rhodamine 123 (Millipore Sigma, Burligton, MA, USA) for 5 seconds, and exciting rhodamine 123 using a white light laser at 511 nm, and a hybrid detector set to 530 nm - 544 nm. The cell wall was visualized using the built-in Differential Interference Contrast (DIC) imaging setting.

***Construction of chloroplasts and mitochondria 3D images.*** The files generated above were exported from the Leica microscope software LasX (version 3.7.4, exported as a .lif file) and converted into .ims files using Imaris File Converter. Whole cell images (in the X, Y, and Z planes) were used to generate 3D reconstructions of chloroplasts and mitochondria in Imaris x64 (version 9.8.0) using the built-in 'Surface' algorithm (set to 0.3 μm surface detail).

***Quantifying chloroplast and mitochondria number and volume.*** After generating 3D reconstructions of mitochondria and chloroplast, described above, the number of cup-shaped and disk-shaped chloroplasts was manually counted. For cup-shaped chloroplasts, orientation of their opening relative to the main vein was also manually counted. For quantifying the volume of cup-shaped and disk-shaped chloroplasts, 3D reconstructions of single chloroplasts were generated as described above, and the volume was calculated by using the built-in Imaris 'Vantage View' tool. A similar approach was used for calculating the total mitochondria volume. For quantifying the volume of mitochondria touching the chloroplast, 3D reconstructions of both organelles within the same cell were first generated, and the mitochondria visibly touching a chloroplast were isolated in Imaris by hand, and the volume of these isolated mitochondria was calculated as described above.

***Cell volume.*** To quantify the total volume of the cell, DIC images of whole cells (i.e. the X,Y planes and all sections of the Z plane that contained the cell) were opened in ImageJ (Version 1.52), and the area of the cell (the area within the cell walls at the median Z-section of a cell, calculated using the built in ImageJ 'Measure' function) was multiplied by the number of Z-sections.

Transmission electron microscopy (TEM)

Leaf disks of 8 mm in diameter (OT206, Owden, Ningbo City, China) were immersed in primary fixative (3-5% glutaraldehyde in 0.1 M cacodylate buffer, pH 6.8-7.1) and incubated for 2 hours at room temperature under vacuum in a fume hood (20 psi). Samples were washed 3 times in 0.1 M cacodylate buffer for 10 minutes each, under vacuum. Samples were post-fixed in

1% osmium tetroxide in 0.1M cacodylate buffer for 2 hours in a vacuum under a fume hood and washed twice 10 minutes each, with 0.1M cacodylate buffer. Gradual dehydration with ethanol was conducted using 50%, 75%, and 95% ethanol in which each step was performed twice for 10 minutes each. Samples were infiltrated with 1:1 95% ethanol:LR white (14381-UC, Electron Microscopy Sciences, Hatfield, PA, USA) for 1 hour at room temperature, then infiltrated with 2 changes of pure LR white for 1 hour (30 minutes for each change) at room temperature, and finally 2 changes of LR white for 1 hour at room temperature (30 minutes for each change). Samples were placed in BEEM capsules (70010-B, Electron Microscopy Sciences, Hatfield, PA, USA) containing fresh LR white and blocks were polymerized at 55ºC for 24 hours.

To confirm the presence of cup-shaped chloroplasts independently, we used another preparation method. Leaf discs of 1 mm diameter were collected by using a biopsy punch (Royaltek, Surgical Design, Lorton, VA, USA) and immediately transferred to 4% paraformaldehyde in phosphate buffer (PBS 1x). For ultrastructural studies, discs were further fixed with 2% glutaraldehyde in 0.1M PIPES, pH 4 overnight, washed and quenched in 50 mM glycine 0.1 M PIPES buffer, pH7.4, before being post-fixed with 2% osmium tetroxide in 0.1M PIPES for 2 hours at room temperature. After osmication, leaf discs were washed with three changes of water, 10 minutes each, dehydrated in a graded ethanol series (15 minutes each in 30%, 50%, 70%, 95% ethanol, and twice in 100% ethanol). Specimens were then infiltrated with Spurr's resin (14300, Electron Microscopy Sciences, Hatfield, PA, USA) starting with 1:1 mixtures of 100% ethanol:Spurr's resin overnight, followed by three changes of pure Spurr's resin over two days and embedded in silicone molds (Product #10535, Ted Pella, Redding, CA, USA).

After polymerization at 60°C for 18–24 hours, ultrathin sections of ~ 90 nm in thickness of the leaf disc were collected on 200 mesh athene thin-bar grids (Ted Pella, Redding, CA, USA) and imaged in a Hitachi 7800 transmission electron microscope at 80 keV (Hitachi, Tokyo, Japan). For each condition and genotype, at least 3 plants were prepared and 3 disks per plant were harvested. The experiment was repeated twice independently. Chemicals were purchased at Millipore Sigma (Burligton, MA, USA) and Electron Microscopy Sciences (Hatfield, PA, USA) and prepared per manufacturer's instructions.

Scanning electron microscopy (SEM)

Leaf disks of 8 mm in diameter were imaged with an Environmental Scanning Electron Microscope (Quanta 200 SEM, Thermofisher, Waltham, MA, USA) equipped with Back Scatter Electron (BSE) detectors in high-vacuum mode. ImageJ was used to evaluate the area occupied by trichomes per leaf area. A threshold was applied to images to separate trichomes of the epidermal surface and the "analyze particles" command was used to measure the area occupied by trichomes.

RCA gene phylogeny

Orthologs of RCA from *T. oblongifolia* and other species were found using the BLAST (version 2.13.0) program tblastn with *Arabidopsis thaliana* RCA (AT2G39730) along with its closest homolog (AT1G73110) protein sequences against a local nucleotide coding sequence database. Primary coding sequences for 25 other genomes were downloaded from Phytozome v.13 (Data S6) (*104*). These were selected based on genome assembly quality and phylogenetic coverage across land plants. For each query gene, BLAST hits were selected up to the other *Arabidopsis* query gene. Matching sequences were translated to amino acids in the first reading frame using the transeq program in the EMBOSS package version 6.6.0 (*117*). Protein alignments were generated with MAFFT version 7.520 using default settings (*118*), and codon alignments were generated from these and the unaligned nucleotide sequences using PAL2NAL version 14 with default settings (*119*). The gene phylogeny was inferred with FastTree version 2.1.11 (*120*) with arguments -nt -gtr -gamma, specifying nucleotide data, the generalized time-reversible substitution model, and a discrete 20-category gamma rate model, respectively. Support values at nodes are local posterior probabilities calculated by FastTree with the Shimodaira-Hasegawa test. A preliminary tree with all matches for both queries exhibited two broadly conserved monophyletic gene clades, each with genes from all species in the database and each with *Sphagnum* and *Selaginella* genes branching from the basal node. This approach ensured that all orthologs had been found for each query gene. Sequences in the RCA gene clade were then realigned separately and the final tree composed of only RCA orthologs was inferred in the same manner, with *Sphagnum* genes set as the outgroup. The tree was visualized using FigTree (http://tree.bio.ed.ac.uk/software/figtree). The associated multiple sequence alignment was generated using Geneious Prime 2023 (www.geneious.com).

Predicted interactors of RCA

From the set of 10817 reciprocal best hit *A. thaliana* homologs in the *T. oblongifolia* genome, predicted RCA-interacting proteins were identified using STRING (version 11.5) (*121*) with default settings.

Graphs and Statistical analysis

Graphs and statistics were performed using GraphPad version 9.0 and R (*122*).

Photos and illustrations

Photos of whole plants were taken with a smartphone and illustrations were created with BioRender.com.

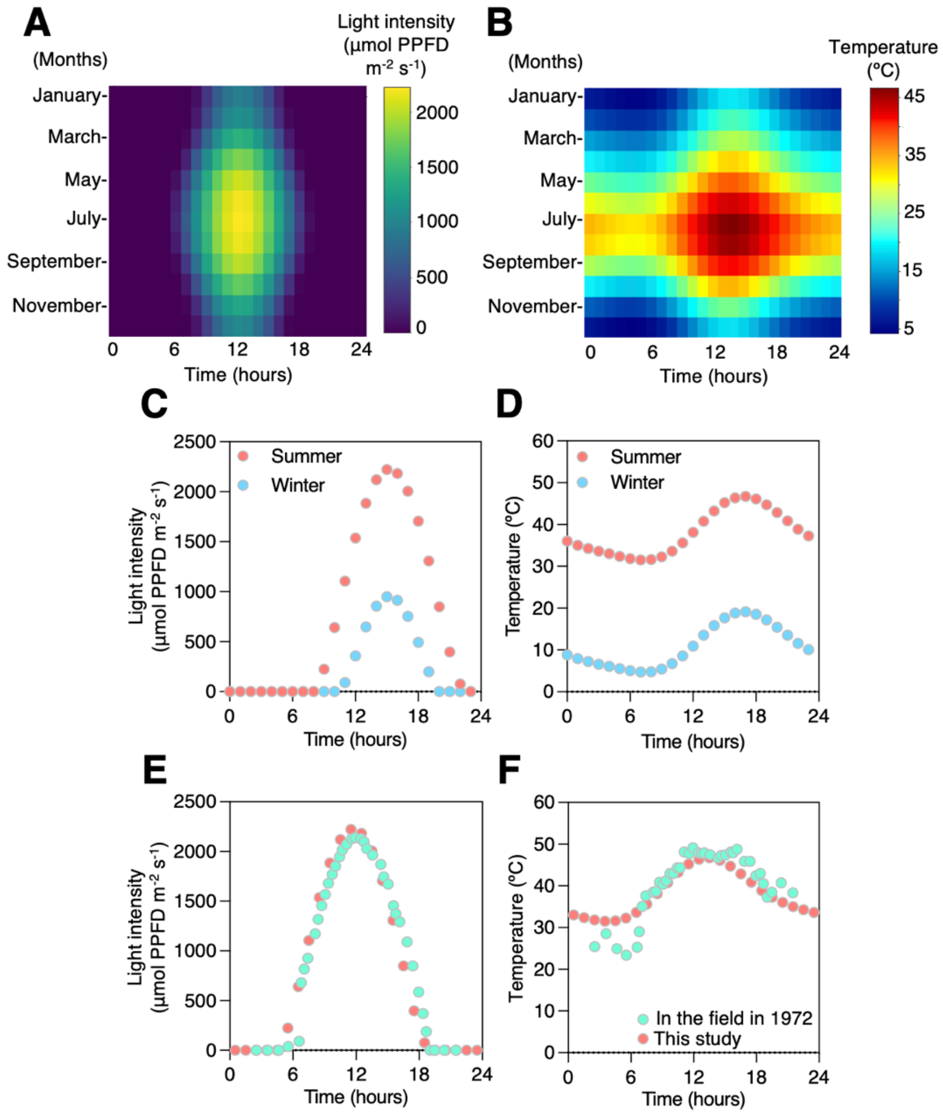


**Fig. S1. Modeling of daily light intensity and temperature at Death Valley, CA.** Modeled daily (**A**) light intensity and **(B)** temperature at Furnace Creek, CA throughout the year according to (*75*). Recreated daily (**C**) light intensity and (**D**) temperature in December (blue) and July (red) at Furnace Creek, CA in programmed custom-built growth chambers. The daylight was shifted by 3 hours from 6 am in the field to 9 am in the custom-built growth chambers. Comparison of modeled (red) and measured (green) in the field (*14*) at Furnace Creek, CA of daily (**E**) light intensity and (**F**) temperature. For comparison, the dawn of custom-built growth chambers was shifted to the corresponding dawn in the field in (E) and (F). Data used to generate the graphs are available in Data S1.

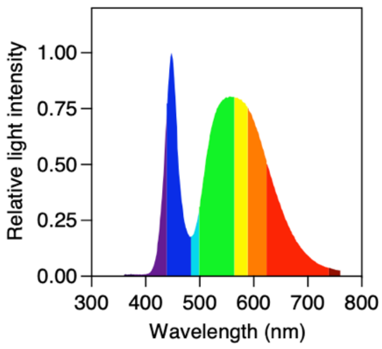


**Fig. S2. Light spectrum in programmed custom-built growth chambers.**
Data used to generate this graph is available in Data S1.

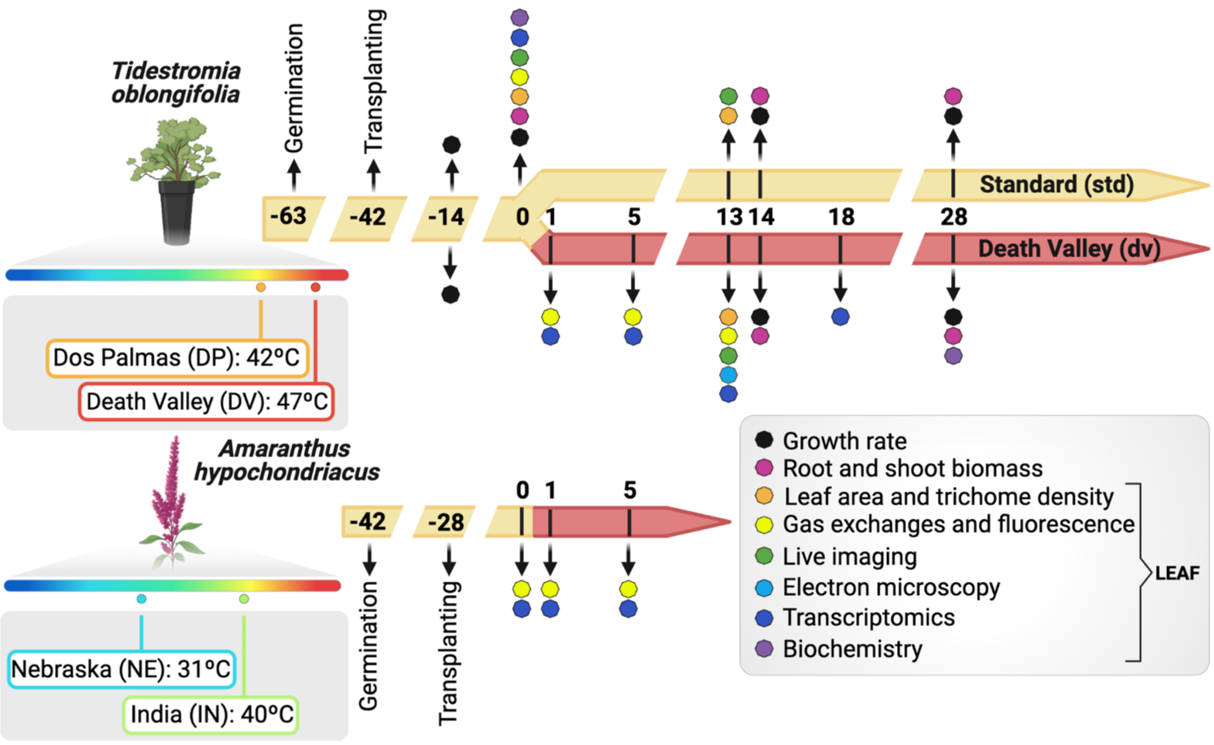


**Fig. S3. Experimental overview and design.** Two accessions of *T. oblongifolia* collected from Death Valley (DV) and Dos Palmas (DP) and two accessions of *A. hypochondriacus* from India (IN) and Nebraska (NE) were selected for this study. Their native regions have a range of annual maximum average temperatures (DV: 47ºC, DP: 42ºC, IN: 40ºC. NE: 35ºC). *T. oblongifolia* plants were grown under standard conditions (std, 31ºC 16h Light / 22ºC 8h Dark) for 9 weeks and transferred to DV summer conditions (dv) for one month or remained under standard conditions (std). Growth rate, root and shoot biomass, leaf area, trichome density, gas-exchange and fluorescence, live imaging, electron microscopy, transcriptomics, and biochemistry were performed with samples harvested on the days indicated in the figure.

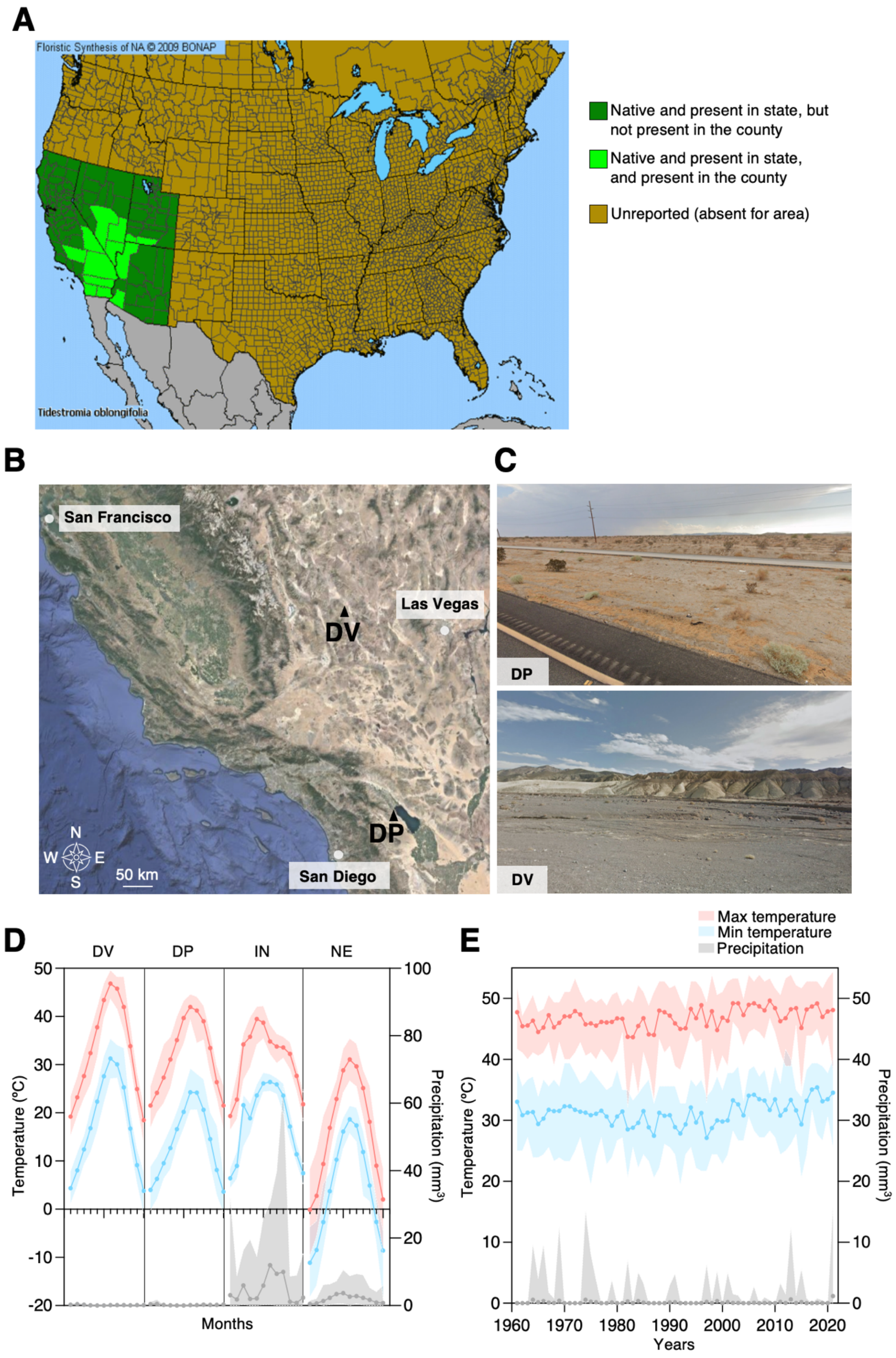
A
Floristic Synthesis of NA © 2009 BONAP
Tidestromia oblongifolia
Native and present in state, but not present in the county
Native and present in state, and present in the county
Unreported (absent for area)
B
San Francisco
Las Vegas
DV
DP
San Diego
N
W
E
S
50 km
C
DP
DV
D
DV
DP
IN
NE
Temperature (°C)
Precipitation (mm³)
Months
E
Max temperature
Min temperature
Precipitation
Temperature (°C)
Precipitation (mm³)
Years

**Fig. S4. Native environment of accessions of *T. oblongifolia* and *A. hypochondriacus*.** (**A**) Distribution of *T. oblongifolia* in states and counties of the US. https://www.pollenlibrary.com/map.aspx?map=Tidestromia-oblongifolia.png (*123*). **(B)** Google map image showing the location of seed collection of Dos Palmas (DP) and Death Valley (DV) accessions. (**C**) Google images captured at 33.322051, -115.996401 (photo taken in September 2021) and at 36.403211, -116.777922 (photo taken in May 2012) where seeds DP (top) and DV (bottom) accessions were collected. (**D**) Native annual climate of accessions in this study from January to December: *T. oblongifolia* DV and DP accessions and *A. hypochondriacus* India (IN) and Nebraska (NE) accessions. Maximum and minimum temperatures are shown in red and blue, respectively and precipitation in gray. Circles and filled areas represent means and range of values for ~70 years. (**E**) Maximum and minimum temperatures and precipitation in July at Death Valley's Furnace Creek Station, CA since 1961. Circles and filled areas represent means and range of values. Data used to generate the graphs are available in Data S1.

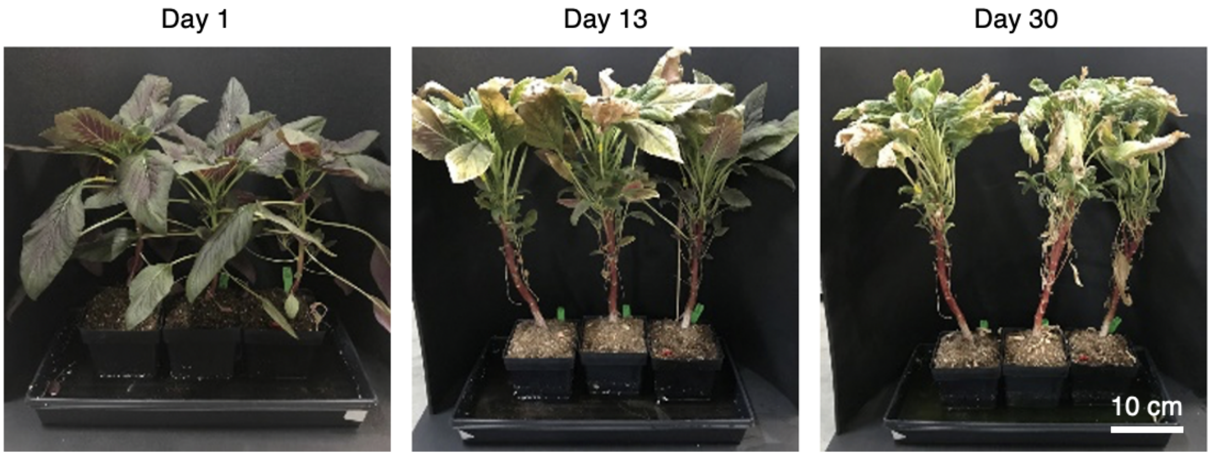


**Fig. S5. Representative images of *A. hypochondriacus* (IN accession) plants grown under DV summer for 1, 13 and 30 days.**

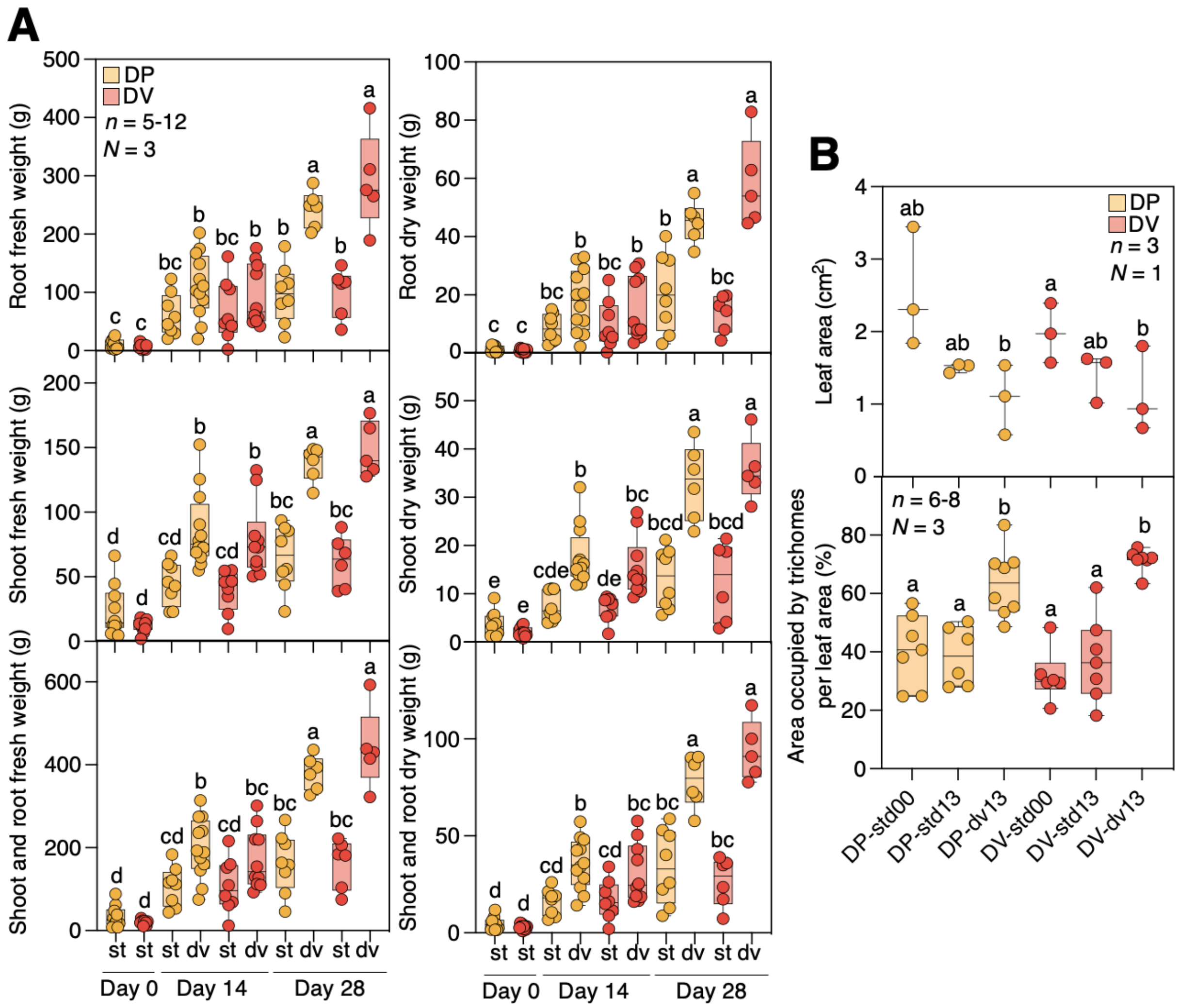


**Fig. S6. Leaf and root characteristics of *T. oblongifolia* in response to DV summer. (A)** Fresh (left panels) and dry (right panels) weights of root (top), shoot (middle), and both (bottom) of *T. oblongifolia* Dos Palmas (DP, orange) and Death Valley (DV, red) accessions harvested under standard conditions before the transfer (day 0), and after growth under DV summer (dv) or corresponding standard conditions (st) for 14 and 28 days. **(B)** Average leaf area per plant and area occupied by trichomes per leaf area for DP and DV accessions grown under standard conditions before the transfer (std00), 13 days under DV summer (dv13) or corresponding standard conditions (std13). Data are presented as box and whisker plots. Different letters indicate statistically different values at $P < 0.05$, as determined by two-way ANOVA followed by a Tukey test. $n$ = number of biological replicates of $N$ independent experiments. Data used to generate the graphs are available in Data S2.

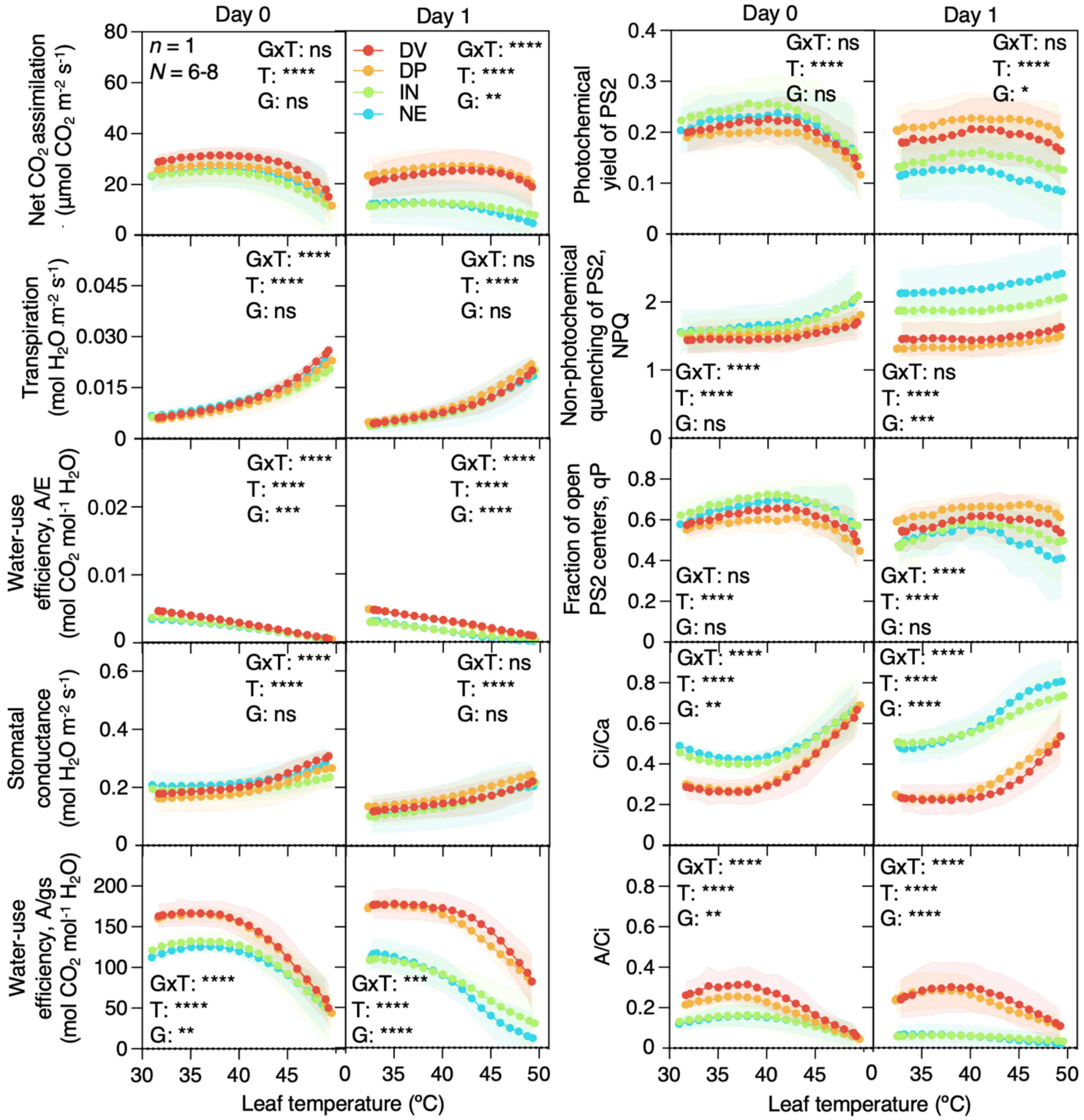


**Fig. S7. Comparison of temperature response curves of photosynthesis parameters between *T. oblongifolia* and *A. hypochondriacus* within a day of being under DV summer.** Temperature response curves of photosynthesis parameters including gas exchange and chlorophyll fluorescence measured with a light intensity of 2200 μmol PPFD $m^{-2}$ $s^{-1}$ and 400 μmol $m^{-1}$ $CO_2$. *T. oblongifolia* (orange: Dos Palmas –DP– accession, red: Death Valley –DV– accession) and *A. hypochondriacus* (green: India –IN– accession, blue: Nebraska –NE– accession) plants were grown under standard conditions before the transfer (Day 0) and transferred to DV summer for 1 day (Day 1). Photosynthetic parameters were monitored using LI-6800. Asterisks indicate statistically significant differences between genotypes (G), temperature (T), or significant interaction between G and T (GxT) using a two-way ANOVA (based on a general linear model) corrected with Geisser-Greenhouse at $P < 0.05$ (*), < 0.01

(**), < 0.001 (***), < 0.0001 (****). Mean ± 95% CI, $n = 1$ leaf per experiment, $N = 6$-8 independent experiments. Data used to generate the graphs are available in Data S3.

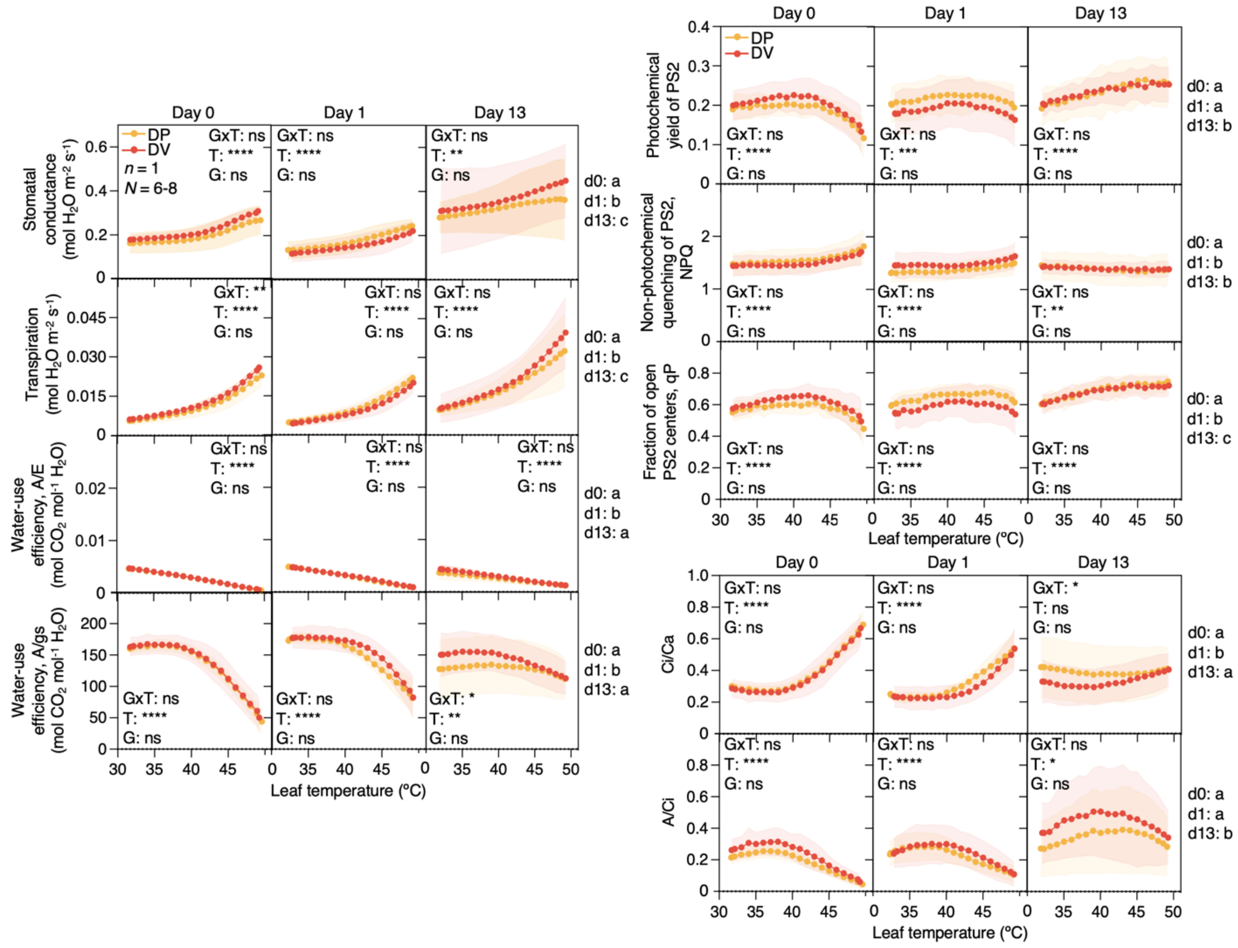


**Fig. S8. Comparison of temperature response curves of photosynthesis parameters in *T. oblongifolia* in days 0, 1, and 13 under DV summer.** Temperature response curves of photosynthesis parameters including gas exchange and chlorophyll fluorescence measured with a light intensity of 2200 µmol PPFD $m^{-2}$ $s^{-1}$ and 400 µmol $m^{-1}$ $CO_2$. *T. oblongifolia* (orange: Dos Palmas –DP– accession, red: Death Valley –DV– accession) plants were grown under standard conditions before the transfer (Day 0) and transferred to DV summer for 1 day (Day 1) and at least 13 days (Day 13). Photosynthesis parameters were monitored using LI-6800. Asterisks indicate statistically significant differences between genotypes (G), temperature (T) or significant interaction between G and T (GxT) using a two-way ANOVA (based on a general linear model) corrected with Geisser-Greenhouse at $P < 0.05$ (*), < 0.01 (**), < 0.001 (***), < 0.0001 (****). Different letters indicate statistically different groups at $P < 0.05$ between days for DV and DP accessions, as determined by three-way ANOVA followed by a Tukey test. Mean ± 95% CI, $n = 1$ leaf per experiment, $N = 6$-8 independent experiments. Data used to generate the graphs are available in Data S3.

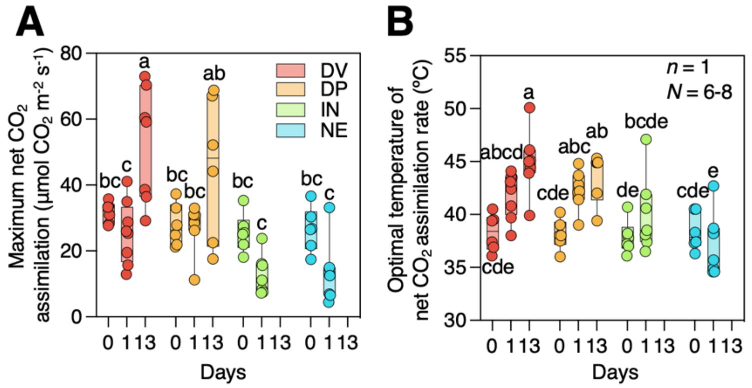


**Fig. S9. Optimal photosynthesis performance in response to DV summer.** Evaluation of (**A**) maximum net $CO_2$ assimilation rate and (**B**) Optimal temperature of net $CO_2$ assimilation rate. Temperature response curves of net $CO_2$ assimilation rate were fitted with a polynomial equation to determine the maximal net $CO_2$ assimilation rate and the temperature at which highest photosynthetic rate was observed (optimal temperature of net $CO_2$ assimilation rate). Results are presented in box and whisker plots. DV (red), DP (orange), IN (green) and NE (blue) accessions were grown under standard conditions (day 0) and transferred to DV summer for 1 day (day 1) or at least 13 days (day 13). Photosynthesis parameters were monitored using LI-6800. Different letters indicate statistically different values at $P < 0.05$, as determined by one-way ANOVA followed by a Tukey test. $n$ = 1 leaf per experiment, $N$ = 6-8 independent experiments. Data used to generate the graphs are available in Data S3.

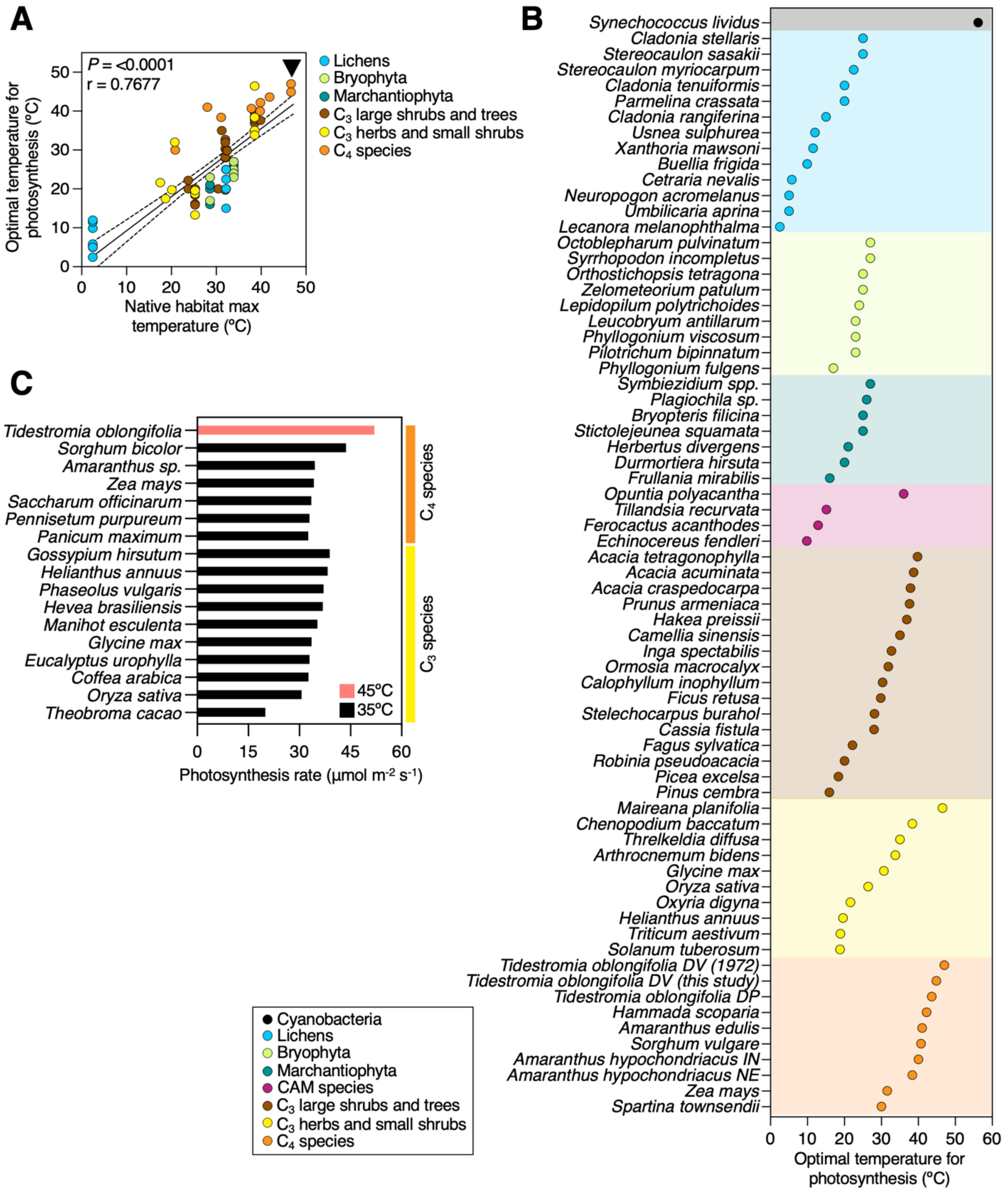


**Fig. S10. Comparison of optimal temperatures of photosynthesis and photosynthesis rates of *T. oblongifolia* under DV summer across plant species. (A)** Correlation between the optimal temperature for photosynthesis and the maximal temperature of the native climates of 73 species grouped in 6 categories: lichens (blue), Bryophyta (light green), Marchantiophyta (dark green), $C_3$ large shrubs and trees (brown), $C_3$ herbs and small shrubs (yellow), and $C_4$ species (orange). 18 published studies were curated (*14*, *23*, *124-139*), and the results of this study were added. A two-tailed Spearman's correlation was computed. The black arrow indicates *T. oblongifolia* measured in the field in the 1970's and in this study. **(B)** Optimal temperature for photosynthesis of various species grouped into categories: cyanobacteria (black), lichens (blue), Bryophyta (light green), Marchantiophyta (dark green), CAM species (magenta), $C_3$ large shrubs and trees (brown), $C_3$ herbs to shrubs (yellow), and $C_4$ species (orange) curated from references (*14*, *23*, *124-139*). Duplicated results for *T. oblongifolia* DV indicate the results obtained in the field in 1972 (14) and the results of this study. **(C)** Maximum photosynthesis rate of *T. oblongifolia* at 45ºC (red) from this study compared to the predicted potential photosynthesis rate of other crops at 35ºC under saturated $CO_2$ and light conditions (black) (*140-141*). For this study, we used plants from the DV accession of *T. oblongifolia* grown for at least 13 days under DV summer. Data used to generate the graphs are available in Data S3.

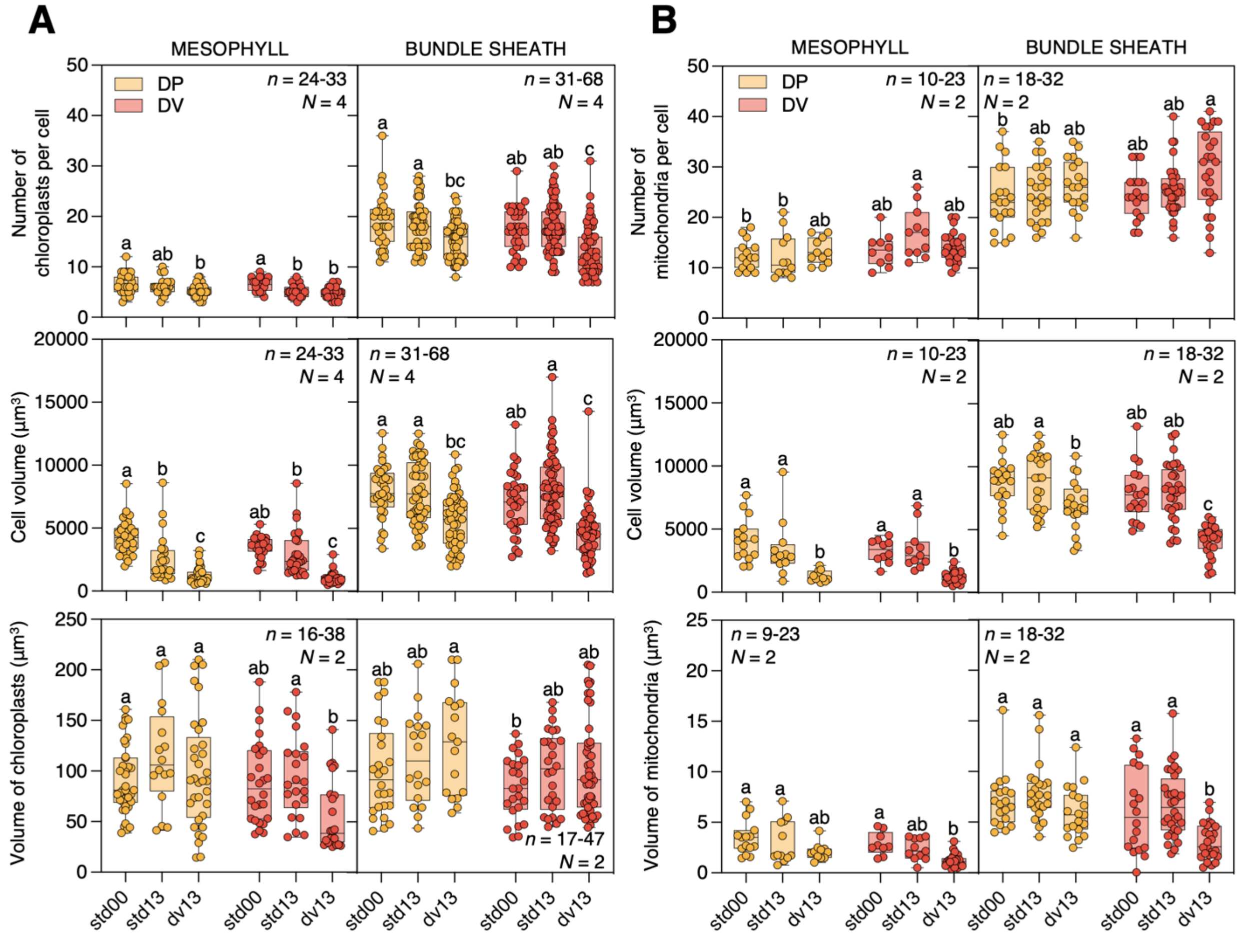


**Fig. S11. Characteristics of cells and organelles in *T. oblongifolia* accessions in response to DV summer by live imaging of mesophyll and bundle sheath cells near tertiary veins.** (**A**) Number of chloroplasts per cell, cell volumes, and volume of chloroplasts in mesophyll (left) and bundle sheath (right) cells. Results are presented as box and whisker plots for Dos Palmas (DP, orange) and Death Valley (DV, red) accessions when grown under standard conditions (std00, std13) and DV summer for 13 days (dv13). Different letters indicate statistically different values at $P < 0.05$, as determined by two-way ANOVA followed by a Tukey test. (**B**) Number of mitochondria per cell, cell volumes, and volume of mitochondria in mesophyll (left) and bundle sheath (right) cells. *n* = biological replicates of *N* independent experiments. Data used to generate the graphs are available in Data S4.

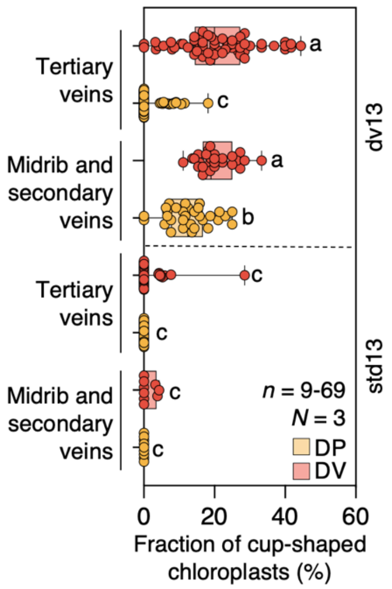


**Fig. S12. Relationship between the order of veins and fraction of cup-shaped chloroplasts in *T. oblongifolia.*** Fraction of cup-shaped chloroplasts in midrib, secondary veins, and tertiary veins in DP (orange) and DV (red) accessions grown for 13 days under DV summer (top) and standard condition (bottom). Different letters indicate statistically different values at $P < 0.05$, as determined by three-way ANOVA followed by a Tukey test. $n$ = biological replicates of $N$ independent experiments. Data used to generate the graphs are available in Data S4.

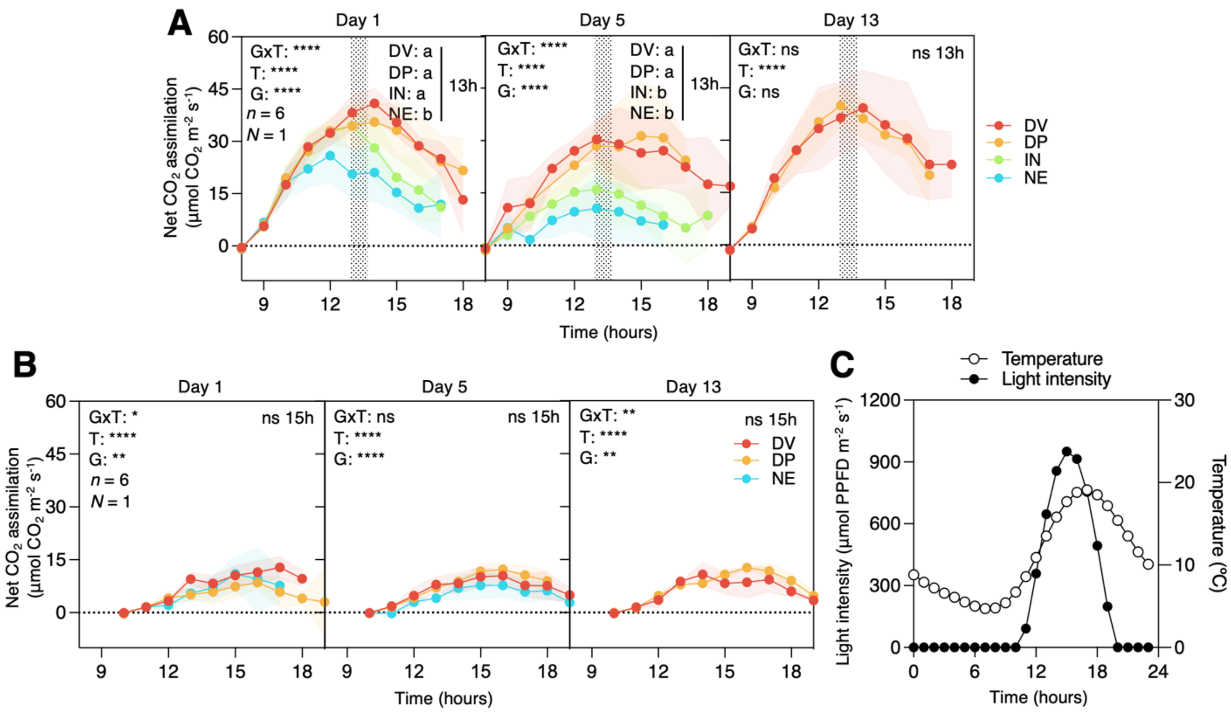


**Fig. S13. Daily photosynthesis performances under DV summer and DV winter.** Daily photosynthesis performances measured under (**A**) DV summer, and (**B**) DV winter. Plants of *T. oblongifolia* (red: DV accession, orange: DP accession) and *A. hypochondriacus* (green: IN accession, blue: NE accession) grown under standard conditions and then transferred to DV summer (top) or DV winter for 1 day (left), 5 days (middle), and 13 days (right). Photosynthesis parameters, including gas exchange and chlorophyll fluorescence, were monitored using the LI-6800. Asterisks indicate statistically significant differences between genotypes (G), time (T), or significant interaction between G and T (GxT) using two-way ANOVA (based on the general linear model) corrected with Geisser-Greenhouse at $P < 0.05$ (*), < 0.01 (**), < 0.001 (***), < 0.0001 (****). This test was followed by False Discovery Rate controlled multiple comparisons at the time indicated. Different letters indicate statistically different values at $P < 0.05$. Mean ± 95% CI, $n = 6$ biological replicates of $N = 1$ independent experiment. Gray areas indicate the time of tissue harvest. (**C**) Daily growth condition of light and temperature under DV winter. Winter natural dawn (8 am) was shifted to 11 am in the custom-built growth chamber. Data used to generate the graphs are available in Data S1 and Data S3.

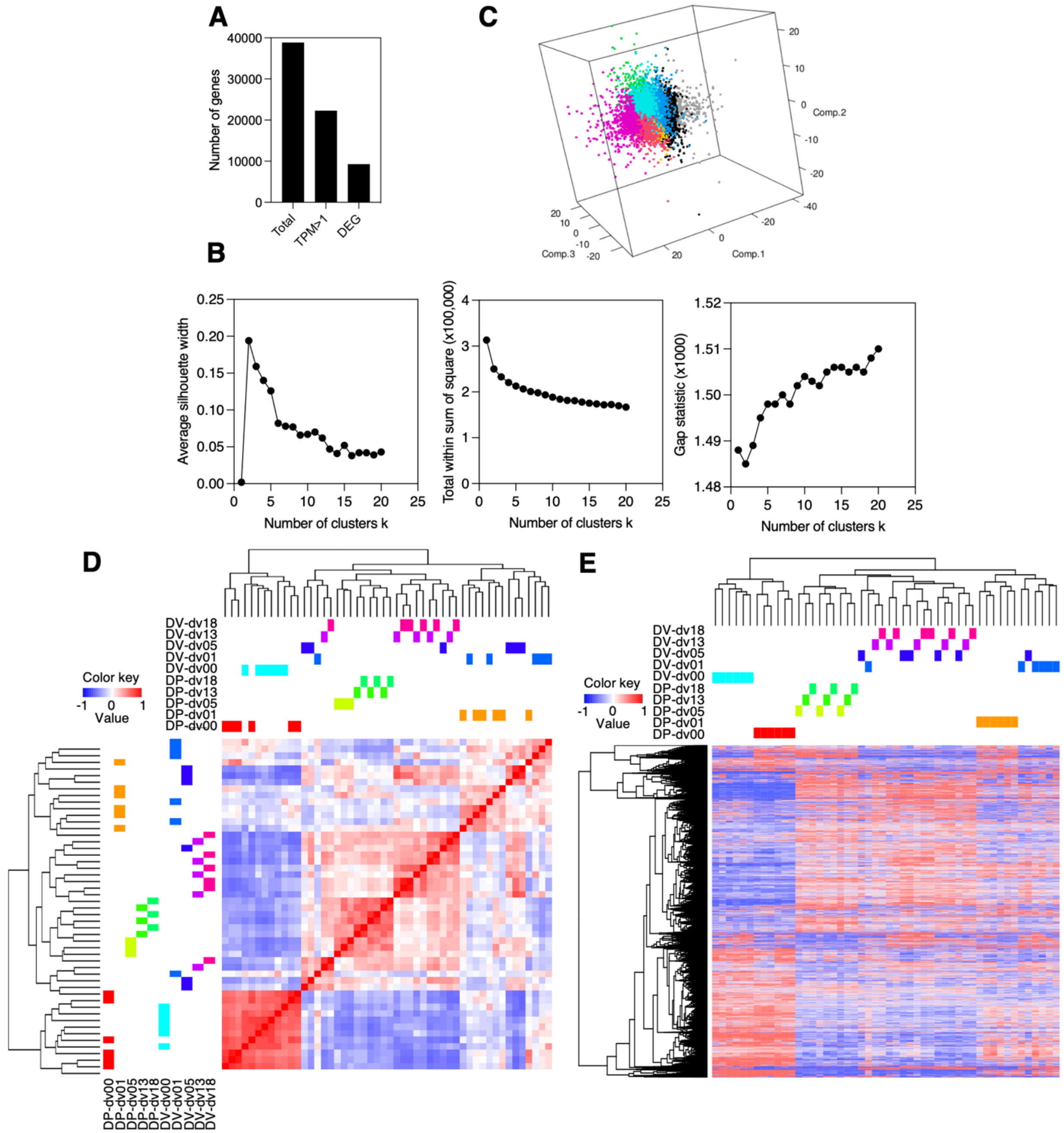


**Fig. S14. Gene expression patterns of samples. (A)** Number of genes in total (38,839), with expression TPM>1 (22,295), and differentially expressed genes (DEG) (9,282). Mitochondrial- and chloroplast-encoded genes were removed from the number of expressed genes with TPM>1 and differentially expressed genes. **(B)** Comparison of methods for determining the number of clusters. The methods considered are Average silhouette method (left), Elbow method (middle), and Gap statistic method (right). **(C)** Principal component analysis of differentially expressed genes fixed with 8 clusters. Heatmaps showing **(D)** correlation of all samples and **(E)** gene expression patterns across the samples. One sample that did not correlate with its replicates was removed. Hierarchical clustering was performed on both dimensions for (D) and (E).

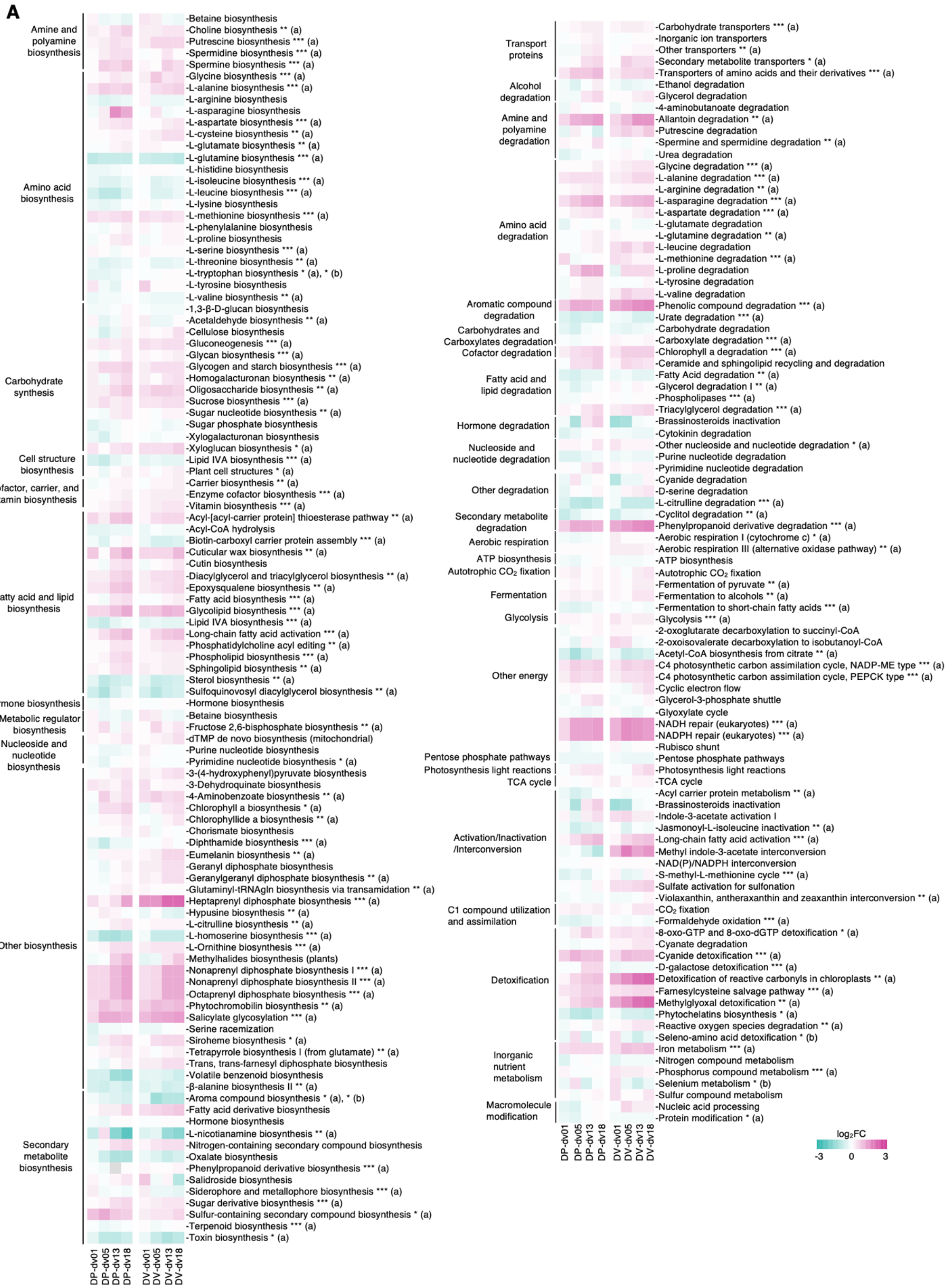
A
Amine and polyamine biosynthesis
-Betaine biosynthesis
-Choline biosynthesis ** (a)
-Putrescine biosynthesis *** (a)
-Spermidine biosynthesis *** (a)
-Spermine biosynthesis *** (a)
Amino acid biosynthesis
-Glycine biosynthesis *** (a)
-L-alanine biosynthesis *** (a)
-L-arginine biosynthesis
-L-asparagine biosynthesis
-L-aspartate biosynthesis *** (a)
-L-cysteine biosynthesis ** (a)
-L-glutamate biosynthesis ** (a)
-L-glutamine biosynthesis *** (a)
-L-histidine biosynthesis
-L-isoleucine biosynthesis *** (a)
-L-leucine biosynthesis *** (a)
-L-lysine biosynthesis
-L-methionine biosynthesis *** (a)
-L-phenylalanine biosynthesis
-L-proline biosynthesis
-L-serine biosynthesis *** (a)
-L-threonine biosynthesis ** (a)
-L-tryptophan biosynthesis * (a), * (b)
-L-tyrosine biosynthesis
-L-valine biosynthesis ** (a)
Carbohydrate synthesis
-1,3-β-D-glucan biosynthesis
-Acetaldehyde biosynthesis ** (a)
-Cellulose biosynthesis
-Gluconeogenesis *** (a)
-Glycan biosynthesis *** (a)
-Glycogen and starch biosynthesis *** (a)
-Homogalacturonan biosynthesis ** (a)
-Oligosaccharide biosynthesis ** (a)
-Sucrose biosynthesis *** (a)
-Sugar nucleotide biosynthesis ** (a)
-Sugar phosphate biosynthesis
-Xylogalacturonan biosynthesis
-Xyloglucan biosynthesis * (a)
Cell structure biosynthesis
-Lipid IVA biosynthesis *** (a)
-Plant cell structures * (a)
Cofactor, carrier, and vitamin biosynthesis
-Carrier biosynthesis ** (a)
-Enzyme cofactor biosynthesis *** (a)
-Vitamin biosynthesis *** (a)
Fatty acid and lipid biosynthesis
-Acyl-[acyl-carrier protein] thioesterase pathway ** (a)
-Acyl-CoA hydrolysis
-Biotin-carboxyl carrier protein assembly *** (a)
-Cuticular wax biosynthesis ** (a)
-Cutin biosynthesis
-Diacylglycerol and triacylglycerol biosynthesis ** (a)
-Epoxysqualene biosynthesis ** (a)
-Fatty acid biosynthesis *** (a)
-Glycolipid biosynthesis *** (a)
-Lipid IVA biosynthesis *** (a)
-Long-chain fatty acid activation *** (a)
-Phosphatidylcholine acyl editing ** (a)
-Phospholipid biosynthesis *** (a)
-Sphingolipid biosynthesis ** (a)
-Sterol biosynthesis ** (a)
-Sulfoquinovosyl diacylglycerol biosynthesis ** (a)
Hormone biosynthesis
-Hormone biosynthesis
Metabolic regulator biosynthesis
-Betaine biosynthesis
-Fructose 2,6-bisphosphate biosynthesis ** (a)
Nucleoside and nucleotide biosynthesis
-dTMP de novo biosynthesis (mitochondrial)
-Purine nucleotide biosynthesis
-Pyrimidine nucleotide biosynthesis * (a)
Other biosynthesis
-3-(4-hydroxyphenyl)pyruvate biosynthesis
-3-Dehydroquinate biosynthesis
-4-Aminobenzoate biosynthesis ** (a)
-Chlorophyll a biosynthesis * (a)
-Chlorophyllide a biosynthesis ** (a)
-Chorismate biosynthesis
-Diphthamide biosynthesis *** (a)
-Eumelanin biosynthesis ** (a)
-Geranyl diphosphate biosynthesis
-Geranylgeranyl diphosphate biosynthesis ** (a)
-Glutaminyl-tRNAgln biosynthesis via transamidation ** (a)
-Heptaprenyl diphosphate biosynthesis *** (a)
-Hypusine biosynthesis ** (a)
-L-citrulline biosynthesis ** (a)
-L-homoserine biosynthesis *** (a)
-L-Ornithine biosynthesis *** (a)
-Methylhalides biosynthesis (plants)
-Nonaprenyl diphosphate biosynthesis I *** (a)
-Nonaprenyl diphosphate biosynthesis II *** (a)
-Octaprenyl diphosphate biosynthesis *** (a)
-Phytochromobilin biosynthesis ** (a)
-Salicylate glycosylation *** (a)
-Serine racemization
-Siroheme biosynthesis * (a)
-Tetrapyrrole biosynthesis I (from glutamate) ** (a)
-Trans, trans-farnesyl diphosphate biosynthesis
-Volatile benzenoid biosynthesis
-β-alanine biosynthesis II ** (a)
Secondary metabolite biosynthesis
-Aroma compound biosynthesis * (a), * (b)
-Fatty acid derivative biosynthesis
-Hormone biosynthesis
-L-nicotianamine biosynthesis ** (a)
-Nitrogen-containing secondary compound biosynthesis
-Oxalate biosynthesis
-Phenylpropanoid derivative biosynthesis *** (a)
-Salidroside biosynthesis
-Siderophore and metallophore biosynthesis *** (a)
-Sugar derivative biosynthesis *** (a)
-Sulfur-containing secondary compound biosynthesis * (a)
-Terpenoid biosynthesis *** (a)
-Toxin biosynthesis * (a)
DP-dv01
DP-dv05
DP-dv13
DP-dv18
DV-dv01
DV-dv05
DV-dv13
DV-dv18
Transport proteins
-Carbohydrate transporters *** (a)
-Inorganic ion transporters
-Other transporters ** (a)
-Secondary metabolite transporters * (a)
-Transporters of amino acids and their derivatives *** (a)
Alcohol degradation
-Ethanol degradation
-Glycerol degradation
Amine and polyamine degradation
-4-aminobutanoate degradation
-Allantoin degradation ** (a)
-Putrescine degradation
-Spermine and spermidine degradation ** (a)
-Urea degradation
Amino acid degradation
-Glycine degradation *** (a)
-L-alanine degradation *** (a)
-L-arginine degradation ** (a)
-L-asparagine degradation *** (a)
-L-aspartate degradation *** (a)
-L-glutamate degradation
-L-glutamine degradation ** (a)
-L-leucine degradation
-L-methionine degradation *** (a)
-L-proline degradation
-L-tyrosine degradation
-L-valine degradation
Aromatic compound degradation
-Phenolic compound degradation *** (a)
-Urate degradation *** (a)
Carbohydrates and Carboxylates degradation
-Carbohydrate degradation
-Carboxylate degradation *** (a)
Cofactor degradation
-Chlorophyll a degradation *** (a)
Fatty acid and lipid degradation
-Ceramide and sphingolipid recycling and degradation
-Fatty Acid degradation ** (a)
-Glycerol degradation I ** (a)
-Phospholipases *** (a)
-Triacylglycerol degradation *** (a)
Hormone degradation
-Brassinosteroids inactivation
-Cytokinin degradation
Nucleoside and nucleotide degradation
-Other nucleoside and nucleotide degradation * (a)
-Purine nucleotide degradation
-Pyrimidine nucleotide degradation
Other degradation
-Cyanide degradation
-D-serine degradation
-L-citrulline degradation *** (a)
-Cyclitol degradation ** (a)
Secondary metabolite degradation
-Phenylpropanoid derivative degradation *** (a)
Aerobic respiration
-Aerobic respiration I (cytochrome c) * (a)
-Aerobic respiration III (alternative oxidase pathway) ** (a)
ATP biosynthesis
-ATP biosynthesis
Autotrophic $CO_2$ fixation
-Autotrophic $CO_2$ fixation
Fermentation
-Fermentation of pyruvate ** (a)
-Fermentation to alcohols ** (a)
-Fermentation to short-chain fatty acids *** (a)
Glycolysis
-Glycolysis *** (a)
Other energy
-2-oxoglutarate decarboxylation to succinyl-CoA
-2-oxoisovalerate decarboxylation to isobutanoyl-CoA
-Acetyl-CoA biosynthesis from citrate ** (a)
-C4 photosynthetic carbon assimilation cycle, NADP-ME type *** (a)
-C4 photosynthetic carbon assimilation cycle, PEPCK type *** (a)
-Cyclic electron flow
-Glycerol-3-phosphate shuttle
-Glyoxylate cycle
-NADH repair (eukaryotes) *** (a)
-NADPH repair (eukaryotes) *** (a)
-Rubisco shunt
Pentose phosphate pathways
-Pentose phosphate pathways
Photosynthesis light reactions
-Photosynthesis light reactions
TCA cycle
-TCA cycle
Activation/Inactivation /Interconversion
-Acyl carrier protein metabolism ** (a)
-Brassinosteroids inactivation
-Indole-3-acetate activation I
-Jasmonoyl-L-isoleucine inactivation ** (a)
-Long-chain fatty acid activation *** (a)
-Methyl indole-3-acetate interconversion
-NAD(P)/NADPH interconversion
-S-methyl-L-methionine cycle *** (a)
-Sulfate activation for sulfonation
-Violaxanthin, antheraxanthin and zeaxanthin interconversion ** (a)
C1 compound utilization and assimilation
-$CO_2$ fixation
-Formaldehyde oxidation *** (a)
Detoxification
-8-oxo-GTP and 8-oxo-dGTP detoxification * (a)
-Cyanate degradation
-Cyanide detoxification *** (a)
-D-galactose detoxification *** (a)
-Detoxification of reactive carbonyls in chloroplasts ** (a)
-Farnesylcysteine salvage pathway *** (a)
-Methylglyoxal detoxification ** (a)
-Phytochelatins biosynthesis * (a)
-Reactive oxygen species degradation ** (a)
-Seleno-amino acid detoxification * (b)
Inorganic nutrient metabolism
-Iron metabolism *** (a)
-Nitrogen compound metabolism
-Phosphorus compound metabolism *** (a)
-Selenium metabolism * (b)
-Sulfur compound metabolism
Macromolecule modification
-Nucleic acid processing
-Protein modification * (a)
DP-dv01
DP-dv05
DP-dv13
DP-dv18
DV-dv01
DV-dv05
DV-dv13
DV-dv18
$log_2FC$
-3
0
3

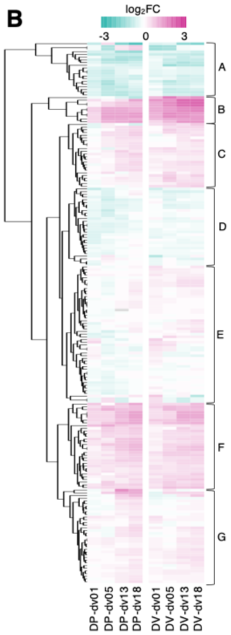


A
-L-nicotianamine biosynthesis ** (a) | Secondary metabolite biosynthesis
-Brassinosteroids inactivation | Hormone degradation
-Brassinosteroids inactivation | Activation/Inactivation/Interconversion
-Volatile Benzenoid biosynthesis | Other biosynthesis
-L-glutamine biosynthesis *** (a) | Amino acid biosynthesis
-Oxalate biosynthesis | Secondary metabolite biosynthesis
-L-homoserine biosynthesis *** (a) | Other biosynthesis
-Toxin biosynthesis * (a) | Secondary metabolite biosynthesis
-Aroma compound biosynthesis * (a), * (b) | Secondary metabolite biosynthesis
-Fatty acid degradation ** (a) | Fatty acid and lipid degradation
-Lipid IVA biosynthesis *** (a) | Cell structure biosynthesis
-Lipid IVA biosynthesis *** (a) | Fatty acid and lipid biosynthesis
-β-alanine biosynthesis II ** (a) | Other biosynthesis
-Urate degradation *** (a) | Aromatic compound degradation
-Sulfoquinovosyl diacylglycerol biosynthesis ** (a) | Fatty acid and lipid biosynthesis
-Acetyl-CoA biosynthesis from citrate ** (a) | Other energy
-L-leucine biosynthesis *** (a) | Amino acid biosynthesis
-Sterol biosynthesis ** (a) | Fatty acid and lipid biosynthesis
-L-citrulline degradation *** (a) | Other degradation
-Phytochelatins biosynthesis * (a) | Detoxification

B
-Methyl indole-3-acetate interconversion | Activation/Inactivation/Interconversion
-Heptaprenyl diphosphate biosynthesis *** (a) | Other biosynthesis
-Detoxification of reactive carbonyls in chloroplasts ** (a) | Detoxification
-Methylglyoxal detoxification ** (a) | Detoxification
-Phenolic compound degradation *** (a) | Aromatic Compound Degradation
-Phenylpropanoid derivative degradation *** (a) | Secondary Metabolite Degradation
-NADH repair (eukaryotes) *** (a) | Other energy
-NADPH repair (eukaryotes) *** (a)| Other energy
-Salicylate glycosylation *** (a) | Other biosynthesis
-Allantoin degradation *** (a) | Amine and Polyamine Degradation

C
-Putrescine degradation | Amine and polyamine degradation
-Siroheme biosynthesis * (a) | Other biosynthesis
-Secondary metabolite transporters * (a) | Transport proteins
-Ceramide and sphingolipid recycling and degradation | Fatty acid and lipid degradation
-Chlorophyll a biosynthesis * (a) | Other biosynthesis
-Spermidine biosynthesis *** (a) | Amine and polyamine biosynthesis
-Photosynthesis light reactions | Photosynthesis light reactions
-L-aspartate biosynthesis *** (a) | Amino acid biosynthesis
-L-aspartate degradation *** (a) | Amino acid degradation
-Indole-3-acetate activation I | Activation/Inactivation/Interconversion
-8-oxo-GTP and 8-oxo-dGTP detoxification * (a) | Detoxification
-Choline biosynthesis ** (a) | Amine and polyamine biosynthesis
-Phospholipid biosynthesis *** (a) | Fatty acid and lipid biosynthesis
-Sphingolipid biosynthesis ** (a) | Fatty acid and lipid biosynthesis
-Triacylglycerol degradation *** (a) | Fatty acid and lipid degradation
-Nitrogen-containing secondary compound biosynthesis | Secondary metabolite biosynthesis
-L-Ornithine biosynthesis *** (a) | Other biosynthesis
-Sugar derivative biosynthesis *** (a) | Secondary metabolite biosynthesis
-Fructose 2,6-bisphosphate biosynthesis ** (a) | Metabolic regulator biosynthesis
-L-leucine degradation | Amino acid degradation
-Glycine biosynthesis *** (a) | Amino acid biosynthesis
-L-valine degradation | Amino acid degradation
-Eumelanin biosynthesis ** (a) | Other biosynthesis
-Sulfate activation for sulfonation | Activation/Inactivation/Interconversion

D
-Cyclitol degradation ** (a) | Secondary metabolite degradation
-Biotin-carboxyl carrier protein assembly *** (a) | Fatty acid and lipid biosynthesis
-Ethanol degradation | Alcohol degradation
-Carbohydrate degradation | Carbohydrates and carboxylates degradation
-Jasmonoyl-L-isoleucine inactivation ** (a) | Activation/Inactivation/Interconversion
-L-arginine biosynthesis | Amino acid biosynthesis
-L-isoleucine biosynthesis *** (a) | Amino acid biosynthesis
-L-threonine biosynthesis ** (a) | Amino acid biosynthesis
-Diphthamide biosynthesis *** (a) | Other biosynthesis
-Xylogalacturonan biosynthesis | Carbohydrate synthesis
-Acyl carrier protein metabolism ** (a) | Activation/Inactivation/Interconversion
-S-methyl-L-methionine cycle *** (a) | Activation/Inactivation/Interconversion
-Pentose phosphate pathways | Pentose phosphate pathways
-Fermentation to Short-Chain Fatty Acids *** (a) | Fermentation
-Formaldehyde Oxidation *** (a) | C1 compound utilization and assimilation
-Acyl-CoA hydrolysis | Fatty acid and lipid biosynthesis
-Purine nucleotide degradation | Nucleoside and nucleotide degradation
-Serine racemization | Other biosynthesis
-Nitrogen compound metabolism | Inorganic Nutrient Metabolism
-Sugar phosphate biosynthesis | Carbohydrate synthesis
-Cytokinin degradation | Hormone degradation
-1,3-β-D-glucan biosynthesis | Carbohydrate synthesis
-L-valine biosynthesis ** (a) | Amino acid biosynthesis
-ATP biosynthesis | ATP biosynthesis
-Protein modification * (a) | Macromolecule modification
-Cyanide degradation | Other degradation
-Hypusine biosynthesis ** (a) | Other biosynthesis
-4-Aminobutanoate degradation | Amine and polyamine degradation
-L-glutamate degradation | Amino acid degradation

F
-Cuticular wax biosynthesis ** (a) | Fatty acid and lipid biosynthesis
-Phytochromobilin biosynthesis ** (a) | Other biosynthesis
-Cyanide detoxification *** (a) | Detoxification
-Octaprenyl diphosphate biosynthesis *** (a) | Other biosynthesis
-Nonaprenyl diphosphate biosynthesis I *** (a) | Other biosynthesis
-Nonaprenyl diphosphate biosynthesis II *** (a) | Other biosynthesis
-Glycolipid biosynthesis *** (a) | Fatty acid and lipid biosynthesis
-L-asparagine degradation *** (a) | Amino acid degradation
-Sulfur-containing secondary compound biosynthesis * (a) | Secondary metabolite biosynthesis
-Spermine biosynthesis *** (a) | Amine and polyamine biosynthesis
-L-alanine biosynthesis *** (a) | Amino acid biosynthesis
-Glycogen and Starch biosynthesis *** (a) | Carbohydrate synthesis
-$C_4$ photosynthetic carbon assimilation cycle, PEPCK type *** (a) | Other energy
-Acyl-[acyl-carrier protein] thioesterase pathway ** (a) | Fatty acid and lipid biosynthesis
-Epoxysqualene biosynthesis ** (a) | Fatty acid and lipid biosynthesis
-Transporters of amino acids and their derivatives *** (a) | Transport proteins
-Long-chain fatty acid activation *** (a) | Fatty acid and lipid biosynthesis
-Long-chain fatty acid activation *** (a) | Activation/Inactivation/Interconversion
-Oligosaccharide biosynthesis ** (a) | Carbohydrate synthesis
-Chlorophyll a degradation *** (a) | Cofactor degradation
-Diacylglycerol and triacylglycerol biosynthesis ** (a) | Fatty acid and lipid biosynthesis
-Farnesylcysteine salvage pathway *** (a) | Detoxification
-L-alanine degradation *** (a) | Amino acid degradation
-L-methionine biosynthesis *** (a) | Amino acid biosynthesis
-Gluconeogenesis *** (a) | Carbohydrate synthesis
-Sucrose biosynthesis *** (a) | Carbohydrate synthesis
-Xyloglucan biosynthesis * (a) | Carbohydrate synthesis
-Putrescine biosynthesis *** (a) | Amine and polyamine biosynthesis
-Fatty acid derivative biosynthesis | Secondary metabolite biosynthesis
-4-aminobenzoate biosynthesis ** (a) | Other biosynthesis
-$C_4$ photosynthetic carbon assimilation cycle, NADP-ME type *** (a) | Other energy
-Iron metabolism *** (a) | Inorganic nutrient metabolism

E
-Sulfur compound metabolism | Inorganic nutrient metabolism
-Phosphorus compound metabolism *** (a) | Inorganic nutrient metabolism
-Aerobic respiration III (alternative oxidase pathway) ** (a) | Aerobic respiration
-Glycolysis *** (a) | Glycolysis
-Cyanate degradation | Detoxification
-Geranylgeranyl diphosphate biosynthesis ** (a) | Other biosynthesis
-Geranyl diphosphate biosynthesis | Other biosynthesis
-Trans, trans-farnesyl diphosphate biosynthesis | Other biosynthesis
-L-phenylalanine biosynthesis | Amino acid biosynthesis
-Glycan biosynthesis *** (a) | Carbohydrate synthesis
-Plant cell structures * (a) | Cell structure biosynthesis
-TCA cycle | TCA cycle
-2-oxoglutarate decarboxylation to succinyl-CoA | Other energy
-L-glutamate biosynthesis ** (a) | Amino acid biosynthesis
-Glutaminyl-tRNAgln biosynthesis via transamidation ** (a) | Other biosynthesis
-L-glutamine degradation ** (a) | Amino acid degradation
-Phenylpropanoid derivative biosynthesis *** (a) | Secondary metabolite biosynthesis
-Carboxylate degradation *** (a) | Carbohydrates and carboxylates degradation
-Phospholipases *** (a) | Fatty acid and lipid degradation
-NAD(P)/NADPH interconversion | Activation/Inactivation/Interconversion
-Cyclic electron flow | Other energy
-D-serine degradation | Other degradation
-Carrier biosynthesis ** (a) | Cofactor, carrier, and vitamin biosynthesis
-Reactive oxygen species degradation ** (a) | Detoxification
-Cellulose biosynthesis | Carbohydrate synthesis
-Urea degradation | Amine and polyamine degradation
-Nucleic acid processing | Macromolecule modification
-L-tyrosine biosynthesis | Amino acid biosynthesis
-L-methionine degradation *** (a) | Amino acid degradation
-2-oxoisovalerate decarboxylation to isobutanoyl-CoA | Other energy
-Betaine biosynthesis | Amine and polyamine biosynthesis
-Betaine biosynthesis | Metabolic regulator biosynthesis
-Acetaldehyde biosynthesis ** (a) | Carbohydrate synthesis
-Siderophore and metallophore biosynthesis *** (a) | Secondary metabolite biosynthesis
-L-serine biosynthesis *** (a) | Amino acid biosynthesis
-Purine nucleotide biosynthesis | Nucleoside and nucleotide biosynthesis
-Rubisco shunt | Other energy
-Glyoxylate cycle | Other energy
-Violaxanthin, antheraxanthin and zeaxanthin interconversion ** (a) | Activation/Inactivation/Interconversion
-Hormone biosynthesis | Hormone biosynthesis
-Hormone biosynthesis | Secondary metabolite biosynthesis
-Pyrimidine nucleotide biosynthesis * (a) | Nucleoside and nucleotide biosynthesis
-Terpenoid biosynthesis *** (a) | Secondary metabolite biosynthesis
-L-tryptophan biosynthesis * (a), * (b) | Amino acid biosynthesis
-L-histidine biosynthesis | Amino acid biosynthesis
-Aerobic respiration I (cytochrome c) * (a) | Aerobic respiration
-L-lysine biosynthesis | Amino acid biosynthesis
-Glycerol degradation I ** (a) | Fatty acid and lipid degradation
-Salidroside biosynthesis | Secondary metabolite biosynthesis
-Seleno-amino acid detoxification * (b) | Detoxification
-Selenium metabolism * (b) | Inorganic nutrient metabolism

G
-L-asparagine biosynthesis | Amino acid biosynthesis
-L-proline degradation | Amino acid degradation
-Methylhalides biosynthesis (plants) | Other biosynthesis
-D-galactose detoxification *** (a) | Detoxification
-L-proline biosynthesis | Amino acid biosynthesis
-Glycerol-3-phosphate shuttle | Other energy
-Sugar nucleotide biosynthesis ** (a) | Carbohydrate synthesis
-Other transporters ** (a) | Transport proteins
-Inorganic ion transporters | Transport proteins
-L-tyrosine degradation | Amino acid degradation
-3-Dehydroquinate biosynthesis | Other biosynthesis
-Other nucleoside and nucleotide degradation * (a) | Nucleoside and nucleotide degradation
-Chorismate biosynthesis | Other biosynthesis
-Spermine and spermidine degradation ** (a) | Amine and polyamine degradation
-Homogalacturonan biosynthesis ** (a) | Carbohydrate synthesis
-Glycerol degradation | Alcohol degradation
-Cutin biosynthesis | Fatty acid and lipid biosynthesis
-Fatty Acid biosynthesis *** (a) | Fatty acid and lipid biosynthesis
-Fermentation to alcohols ** (a) | Fermentation
-$CO_2$ fixation | C1 compound utilization and assimilation
-Autotrophic $CO_2$ fixation | Autotrophic $CO_2$ fixation
-Fermentation of pyruvate ** (a) | Fermentation
-3-(4-hydroxyphenyl)pyruvate biosynthesis | Other biosynthesis
-Vitamin biosynthesis *** (a) | Cofactor, carrier, and vitamin biosynthesis
-Phosphatidylcholine acyl editing ** (a) | Fatty acid and lipid biosynthesis
-L-cysteine biosynthesis ** (a) | Amino acid biosynthesis
-dTMP de novo biosynthesis (mitochondrial) | Nucleoside and nucleotide biosynthesis
-Glycine degradation *** (a) | Amino acid degradation
-Chlorophyllide a biosynthesis ** (a) | Other biosynthesis
-Tetrapyrrole biosynthesis I (from glutamate) ** (a) | Other biosynthesis
-Enzyme cofactor biosynthesis *** (a) | Cofactor, carrier, and vitamin biosynthesis
-Carbohydrate transporters *** (a) | Transport proteins
-L-arginine degradation ** (a) | Amino acid degradation
-L-citrulline biosynthesis ** (a) | Other biosynthesis
-Pyrimidine nucleotide degradation | Nucleoside and nucleotide degradation

**Fig. S15. TidestromiaCyc v1.0.0 dashboard of regulated pathways in response to DV summer in *T. oblongifolia.*** A heat map displaying the $\log_2$ fold change ($\log_2$FC) of gene expression of expressed metabolic enzymes (TPM>1) under DV summer relative to standard condition (**A,B**). Pathways were hierarchically clustered in (B). Mitochondrial- and chloroplast-encoded genes were not considered. Asterisks indicate statistically significant differences between conditions (a) or genotypes (b) using three-way ANOVA at $P < 0.05$ (*), < 0.01 (**), < 0.001 (***). *P* values were corrected for multiple comparisons using the False Discovery Rate (FDR). The details of pathways and their superclass pathways in *T. oblongifolia* are shown on the right in black and gray, respectively. Data used to generate the graphs are available in Data S5.

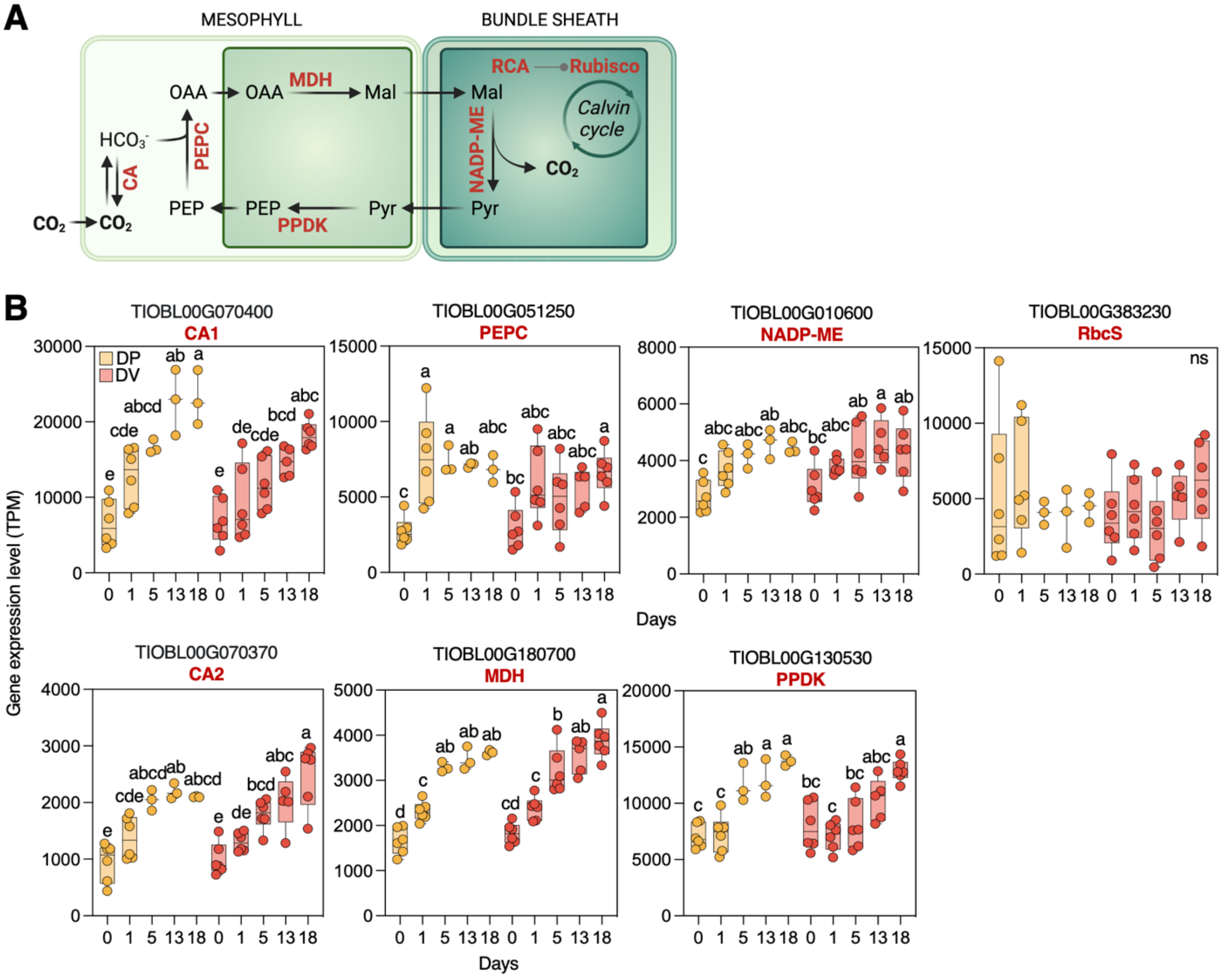


**Fig. S16. Gene expression of C4 photosynthesis pathway of *T. oblongifolia.*** **(A)** A schematic diagram illustrating C4-NADP-ME pathway of *T. oblongifolia* occurring in mesophyll (light green) and bundle sheath (dark green) cells. The chloroplast of each cell is presented in darker colors compared to cytoplasm. Proteins in each reaction are shown in red. **(B)** Gene expression in TPM (Transcripts Per kilobase Million) of key proteins and regulators of NADP-ME C4 pathways of *T. oblongifolia* Dos Palmas (DP, orange) and Death Valley (DV, red) accessions grown under DV summer. Different letters indicate statistically different groups at $P < 0.05$, as determined by two-way ANOVA followed by a Tukey test. CA = β-carbonic anhydrase, PEPC = phosphoenolpyruvate carboxylase, MDH = malate dehydrogenase (NADP), NADP-ME = NADP-dependent malic enzyme, PPDK = pyruvate phosphate dikinase, Rubisco = ribulose-1,5-bisphosphate carboxylase-oxygenase, RCA = rubisco activase. $n$ = 3-6 biological replicates of $N$ = 2 independent experiments.

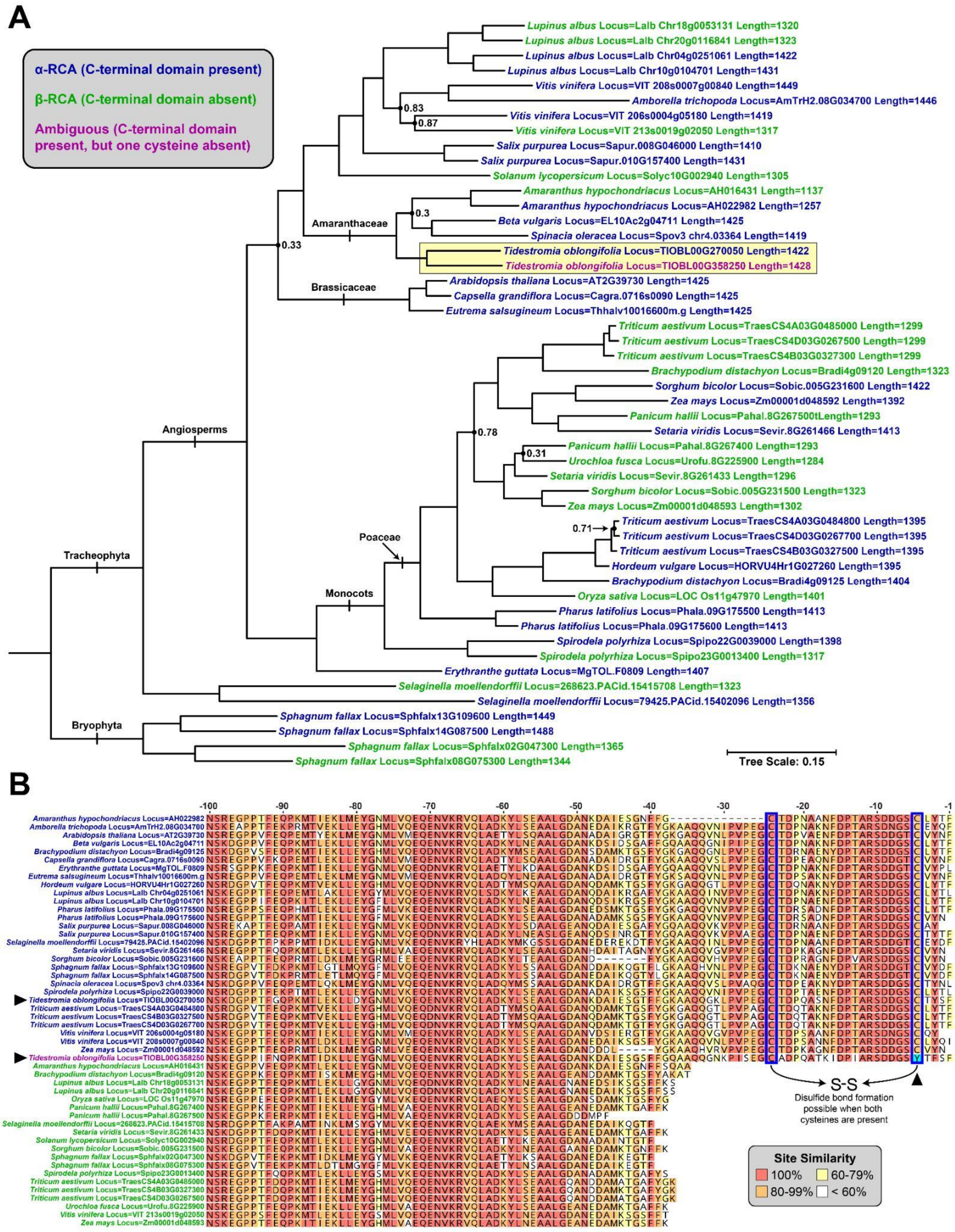
A
α-RCA (C-terminal domain present)
β-RCA (C-terminal domain absent)
Ambiguous (C-terminal domain present, but one cysteine absent)
Lupinus albus Locus=Lalb Chr18g0053131 Length=1320
Lupinus albus Locus=Lalb Chr20g0116841 Length=1323
Lupinus albus Locus=Lalb Chr04g0251061 Length=1422
Lupinus albus Locus=Lalb Chr10g0104701 Length=1431
Vitis vinifera Locus=VIT 208s0007g00840 Length=1449
Amborella trichopoda Locus=AmTrH2.08G034700 Length=1446
Vitis vinifera Locus=VIT 206s0004g05180 Length=1419
Vitis vinifera Locus=VIT 213s0019g02050 Length=1317
Salix purpurea Locus=Sapur.008G046000 Length=1410
Salix purpurea Locus=Sapur.010G157400 Length=1431
Solanum lycopersicum Locus=Solyc10G002940 Length=1305
Amaranthus hypochondriacus Locus=AH016431 Length=1137
Amaranthus hypochondriacus Locus=AH022982 Length=1257
Beta vulgaris Locus=EL10Ac2g04711 Length=1425
Spinacia oleracea Locus=Spov3 chr4.03364 Length=1419
Tidestromia oblongifolia Locus=TIOBL00G270050 Length=1422
Tidestromia oblongifolia Locus=TIOBL00G358250 Length=1428
Arabidopsis thaliana Locus=AT2G39730 Length=1425
Capsella grandiflora Locus=Cagra.0716s0090 Length=1425
Eutrema salsugineum Locus=Thhalv10016600m.g Length=1425
Triticum aestivum Locus=TraesCS4A03G0485000 Length=1299
Triticum aestivum Locus=TraesCS4D03G0267500 Length=1299
Triticum aestivum Locus=TraesCS4B03G0327300 Length=1299
Brachypodium distachyon Locus=Bradi4g09120 Length=1323
Sorghum bicolor Locus=Sobic.005G231600 Length=1422
Zea mays Locus=Zm00001d048592 Length=1392
Panicum hallii Locus=Pahal.8G267500tLength=1293
Setaria viridis Locus=Sevir.8G261466 Length=1413
Panicum hallii Locus=Pahal.8G267400 Length=1293
Urochloa fusca Locus=Urofu.8G225900 Length=1284
Setaria viridis Locus=Sevir.8G261433 Length=1296
Sorghum bicolor Locus=Sobic.005G231500 Length=1323
Zea mays Locus=Zm00001d048593 Length=1302
Triticum aestivum Locus=TraesCS4A03G0484800 Length=1395
Triticum aestivum Locus=TraesCS4D03G0267700 Length=1395
Triticum aestivum Locus=TraesCS4B03G0327500 Length=1395
Hordeum vulgare Locus=HORVU4Hr1G027260 Length=1395
Brachypodium distachyon Locus=Bradi4g09125 Length=1404
Oryza sativa Locus=LOC Os11g47970 Length=1401
Pharus latifolius Locus=Phala.09G175500 Length=1413
Pharus latifolius Locus=Phala.09G175600 Length=1413
Spirodela polyrhiza Locus=Spipo22G0039000 Length=1398
Spirodela polyrhiza Locus=Spipo23G0013400 Length=1317
Erythranthe guttata Locus=MgTOL.F0809 Length=1407
Selaginella moellendorffii Locus=268623.PACid.15415708 Length=1323
Selaginella moellendorffii Locus=79425.PACid.15402096 Length=1356
Sphagnum fallax Locus=Sphfalx13G109600 Length=1449
Sphagnum fallax Locus=Sphfalx14G087500 Length=1488
Sphagnum fallax Locus=Sphfalx02G047300 Length=1365
Sphagnum fallax Locus=Sphfalx08G075300 Length=1344
0.83
0.87
0.3
0.33
0.78
0.31
0.71
Amaranthaceae
Brassicaceae
Angiosperms
Tracheophyta
Poaceae
Monocots
Bryophyta
Tree Scale: 0.15
B
-100 -90 -80 -70 -60 -50 -40 -30 -20 -10 -1
S-S
Disulfide bond formation possible when both cysteines are present
Site Similarity
100%
60-79%
80-99%
< 60%

**Fig. S17. Gene phylogeny and amino acid alignment of RCA for *T. oblongifolia* and 25 reference genomes. (A)** Gene phylogeny of RCA for *T. oblongifolia* and 25 reference genomes. Node support values are local posterior probabilities (LPP) calculated by FastTree with the Shimodaira-Hasegawa test and are omitted when ≥ 0.9. Gene names are colored to indicate presence/absence of the α-RCA C-terminal domain. The ambiguous *T. oblongifolia* gene copy which lacks one of the conserved cysteine residues is denoted with purple. Even with the inclusion of three other Amaranthaceae genomes, the two RCA sequences from *T. oblongifolia* (yellow box) are each other's closest relative, indicating a recent and possibly *Tidestromia*-specific duplication event. The putative α and β isoforms of RCA do not form monophyletic gene clades, but rather appear to have evolved multiple times convergently in diverse clades. **(B)** Amino acid alignment of the RCA C-terminal region of all sequences in the RCA gene phylogeny. The last 100 amino acid sites of the alignment are shown. Sequences are grouped as putative α (top, blue sequence IDs) and β (bottom, green sequence IDs) isoforms, with the ambiguous *T. oblongifolia* sequence denoted in purple. Blue boxes around sites -24 and -5 denote the two redox-sensitive cysteine residues. Black arrowheads denote the two *T. oblongifolia* genes (left) and the cysteine-to-tyrosine substitution in TIOBL00G358250 (right). The alignment color scale indicates site similarity under the Blosum62 substitution scoring matrix. Data used to generate the graphs are available in Data S6.

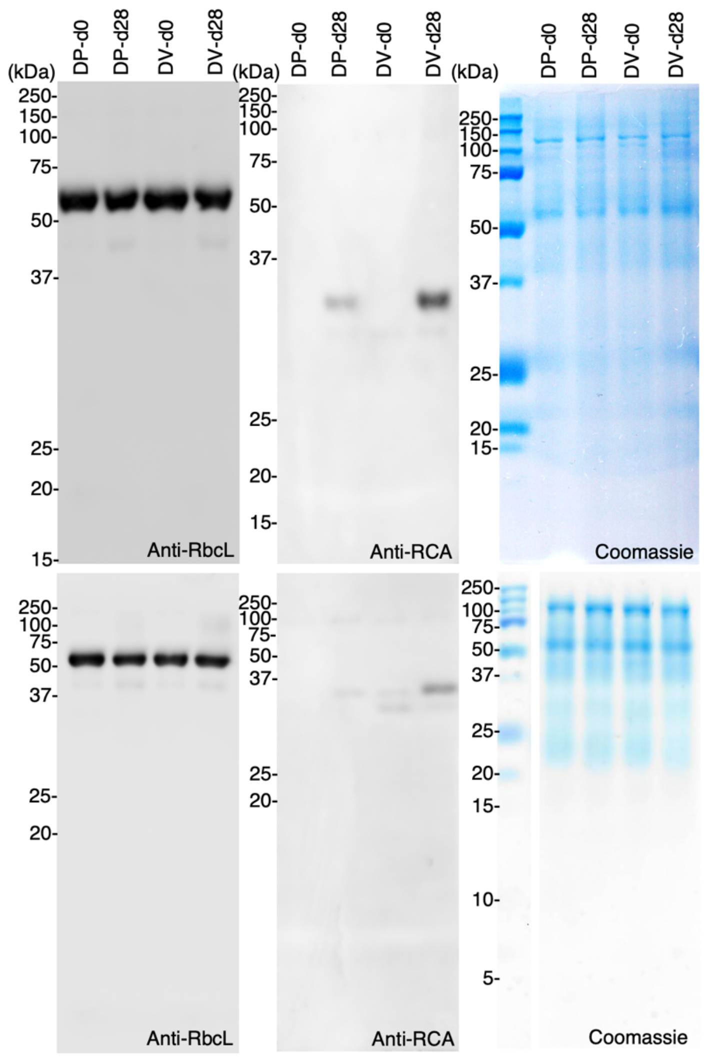


**Fig. S18. Western blot using RbcL and RCA antibodies.** Leaf proteins of Dos Palmas (DP) accessions and Death Valley (DV) grown under standard conditions or under DV summer for 28 days, analyzed by Western blot using RbcL (Rubisco Large unit) and RCA (Rubisco activase) antibodies. Two independent biological experiments with indication of molecular weight marks on the left.

**Table S1. Summary of genome sequencing data for *T. oblongifolia*.**

<table>
<tr><th colspan="4">Genome sequencing (Gb)</th><th colspan="6">Genome size estimation by 21-mer analysis</th></tr>
<tr><td>DNB</td><td>PacBio</td><td>PacBio Type</td><td>mRNA-seq</td><td>K-mer size</td><td>K-mer number (Gb)</td><td>Peak depth (×)</td><td>Estimated genome size (Gb)</td><td colspan="2">Heterozygous ratio (%)</td></tr>
<tr><td>155.41</td><td>158.34</td><td>CLR</td><td>50.4</td><td>21</td><td>82.79</td><td>38</td><td>2.17</td><td colspan="2">0.57</td></tr>
<tr><th colspan="4">PacBio and assembled genome size</th><th colspan="6">Completeness evaluation of genome with BUSCO database and mapping ratio of short reads and transcriptome data</th></tr>
<tr><td>Bases (Gb)</td><td>Reads (M)</td><td>Depth (×)</td><td>Assembled Genome size (Gb)</td><td colspan="2">BUSCO value</td><td colspan="2">Mapping ratio of DNB-seq</td><td colspan="2">Mapping ratio of mRNA-seq</td></tr>
<tr><td>158.34</td><td>6.65</td><td>72.97</td><td>2.16</td><td colspan="2">97.8</td><td colspan="2">99.85</td><td colspan="2">95.62</td></tr>
<tr><th colspan="10">The contig N value of assembled genome (Mb)</th></tr>
<tr><td>0</td><td>10</td><td>20</td><td>30</td><td>40</td><td>50</td><td>60</td><td>70</td><td>80</td><td>90</td></tr>
<tr><td>32.8</td><td>19.76</td><td>16.54</td><td>14.23</td><td>10.75</td><td>9.19</td><td>7.72</td><td>6.06</td><td>3.89</td><td>2.07</td></tr>
</table>

**Table S2. Repeat annotations of *T. oblongifolia* genome.**

| Class | | Count | Masked (bp) | %masked |
|---|---|---|---|---|
| DNA | DTA | 189504 | 33209965 | 1.54 |
| | DTC | 128101 | 32528386 | 1.51 |
| | DTH | 44514 | 10184373 | 0.47 |
| | DTM | 178508 | 38260207 | 1.77 |
| | DTT | 1331392 | 183759010 | 8.52 |
| | Helitron | 779114 | 158629713 | 7.35 |
| LINE | unknown | 1563 | 851140 | 0.04 |
| LTR | Copia | 257556 | 176047013 | 8.16 |
| | Gypsy | 899683 | 765910641 | 35.50 |
| | unknown | 1149531 | 443846002 | 20.57 |
| MITE | DTA | 27847 | 4066546 | 0.19 |
| | DTC | 9083 | 1296846 | 0.06 |
| | DTH | 8033 | 1108706 | 0.05 |
| | DTM | 45517 | 6351939 | 0.29 |
| | DTT | 138778 | 10117133 | 0.47 |
| TIR | EnSpm_CACTA | 356 | 39659 | 0.00 |
| | MuDR_Mutator | 1571 | 820950 | 0.04 |
| | PIF_Harbinger | 137 | 69987 | 0.00 |
| | hAT | 75 | 48588 | 0.00 |
| Unknown | | 112371 | 26995496 | 1.25 |
| pararetrovirus | | 431 | 279737 | 0.01 |
| total interspersed | | 5303665 | 1894422037 | 87.80 |
| Low_complexity | | 16751 | 827053 | 0.04 |
| Simple_repeat | | 244964 | 34302916 | 1.59 |
| Total | | 5565380 | 1929552006 | 89.43 |

**Table S3. List of softwares used in this study.**

| Analysis | Software | Version | Parameter | Notes |
|---|---|---|---|---|
| Genome size estimation | gce | 1.0.0 | -k 21 -a 0 -d 0 for kmer_freq_hash and -m 1 -b 1 for gce | |
| Genome assembly | Canu | 2.0 | genomeSize = 2.17g, minOverlapLength = 700, minReadLength = 1000 | |
| Genome assembly | minimap2 | 2.1 | -x asm5 and purge_dups (v1.2.3) with -T 2 | |
| Genome assembly | Pilon | 1.24 | | |
| Genome repeat identification | EDTA | 1.9.6 | --anno 1 --force 1 --debug 1 --sensitive 1 --evaluate 1 | |
| Genome repeat identification | RepeatMasker | 4.1.2 | -a -html -gff | http://repeatmasker.org |
| Gene model prediction | Maker | 3.01.03 | default parameter | https://www.yandell-lab.org/software/maker.html |
| Gene model prediction | Trinity | 2.11 | default parameter | |
| Gene model prediction | Cufflinks | 2.2.1 | --library-type fr-unstranded | |
| Gene model prediction | StringTie | 2.2.1 | default parameter | |
| Gene model prediction | Mikado | 2.3.3 | default parameter | |
| Gene model prediction | PASA | 2.5.2 | default parameter | |
| Gene model prediction | Fgenesh++ | 7.0 | default parameter | http://www.softberry.com/ |
| Gene model prediction | Augustus | 3.5.0 | default parameter | |
| Gene model prediction | GeneMark-ET | 4.71_lic | default parameter | |
| Gene model prediction | Braker2 | 2.1.6 | --softmasking --gff3 --nocleanup | |
| Redundancy Removal | CD-HIT | 4.8.1 | -c 0.99 -G 0 and -aS 0.75 | |
| Functional Annotation | BLAST+ | 2.10.0 | Nt, selected species: -evalue 1e-5 | |
| Functional Annotation | DIAMOND | 2.0.4 | NR, Swiss-Prot: --evalue 1e-5; KOG/COG: --evalue = 1e-3 | http://www.ncbi.nlm.nih.gov/KOG/, https://www.ebi.ac.uk/uniprot/ |
| Functional Annotation | E2P2 | 4.0 | | Enzyme annotation, https://github.com/carnegie/E2P2 |

| | | | | |
|---|---|---|---|---|
| Functional Annotation | InterProScan | 5.38 | | GO annotation, http://www.geneontology.org/ |
| Functional Annotation | Pathway Tools | 24.0 | | SRI International |
| Functional Annotation | SAVI | 3.1 | | https://plantcyc.org/downloads?qt-downloads=1 |
| Differential Expression Analysis | DESeq2 | 1.26.0 | | |
| Differential Expression Analysis | HISAT2 | 2.2.1 | --score-min L,0,-0.875 | |
| Differential Expression Analysis | R | 3.6.2 | | Kmeans |
| Genome Assembly and Annotation Completeness | BUSCO | 4.0.5 | | |
| Orthologs | OrthoFinder | 2.3.12 | default parameter | |
| Gene ontology enrichment | Cytoscape along with its plugin Bingo version 3.0.3 | 3.9.1 | | http://www.cytoscape.org/ |
| Prediction of interactants | STRING | 11.5 | | https://string-db.org/ |

**Data S1. Climate dataset**

**Data S2. Leaf and root characteristics dataset**

**Data S3. Photosynthesis measurement dataset**

**Data S4. Microscopy dataset**

**Data S5. Transcriptomics dataset**

**Data S6. Specific genes dataset**
